# A molecular filter for the cnidarian stinging response

**DOI:** 10.1101/2020.04.04.025338

**Authors:** Keiko Weir, Christophe Dupre, Lena van Giesen, Amy S.Y. Lee, Nicholas W. Bellono

## Abstract

All animals detect and integrate diverse environmental signals to mediate behavior. Cnidarians, including jellyfish and sea anemones, both detect and capture prey using stinging cells called nematocytes which fire a venom-covered barb via an unknown triggering mechanism. Here, we show that nematocytes from *Nematostella vectensis* use a specialized voltage-gated calcium channel (nCa_v_) to distinguish salient sensory cues and control the explosive discharge response. Adaptations in nCa_v_ confer unusually-sensitive, voltage-dependent inactivation to inhibit responses to non-prey signals, such as mechanical water turbulence. Prey-derived chemosensory signals are synaptically transmitted to acutely relieve nCa_v_ inactivation, enabling mechanosensitive-triggered predatory attack. These findings reveal a molecular basis for the cnidarian stinging response and highlight general principles by which single proteins integrate diverse signals to elicit discrete animal behaviors.

## Introduction

Jellyfish, sea anemones, and hydrozoans of the Cnidarian phylum use specialized cells called cnidocytes to facilitate both sensation and secretion required for prey capture and defense (Watson and Mire-Thibodeaux, 1994). Two major types of cnidocytes contribute to prey capture by the tentacles of the starlet sea anemone *(Nematostella vectensis*, **Fig. 1A**): (1) spirocytes, anthozoan-specific cells that extrude a thread-like organelle to ensnare prey, and (2) nematocytes, pan-cnidarian cells which eject a single-use venom-covered barb to mediate stinging (Babonis and Martindale, 2017). Sensory cues from prey act on nematocytes to trigger the explosive discharge of a specialized organelle (nematocyst) at an acceleration of up to 5.41×10^6^ g, among the fastest of any biological process (Holstein and Tardent, 1984, Nuchter et al., 2006) (**Fig. 1B**). The nematocyst can only be discharged once and therefore stinging represents an energetically expensive process that is likely tightly regulated (Watson and Mire-Thibodeaux, 1994, Babonis and Martindale, 2014). Indeed, simultaneously presented chemical and mechanical (chemo-tactile) cues are required to elicit nematocyte discharge (Pantin, 1942b, Watson and Hessinger, 1989, Watson and Hessinger, 1992, Anderson and Bouchard, 2009). Electrical stimulation of nematocytes increases the probability of discharge in a calcium (Ca^2+^)-dependent manner (Anderson and Mckay, 1987, Mckay and Anderson, 1988, Santoro and Salleo, 1991, Gitter et al., 1994, Watson and Hessinger, 1994, Anderson and Bouchard, 2009), but direct recordings from nematocytes are limited and thus mechanisms by which environmental signals control discharge are not well studied.

**Figure 1:**
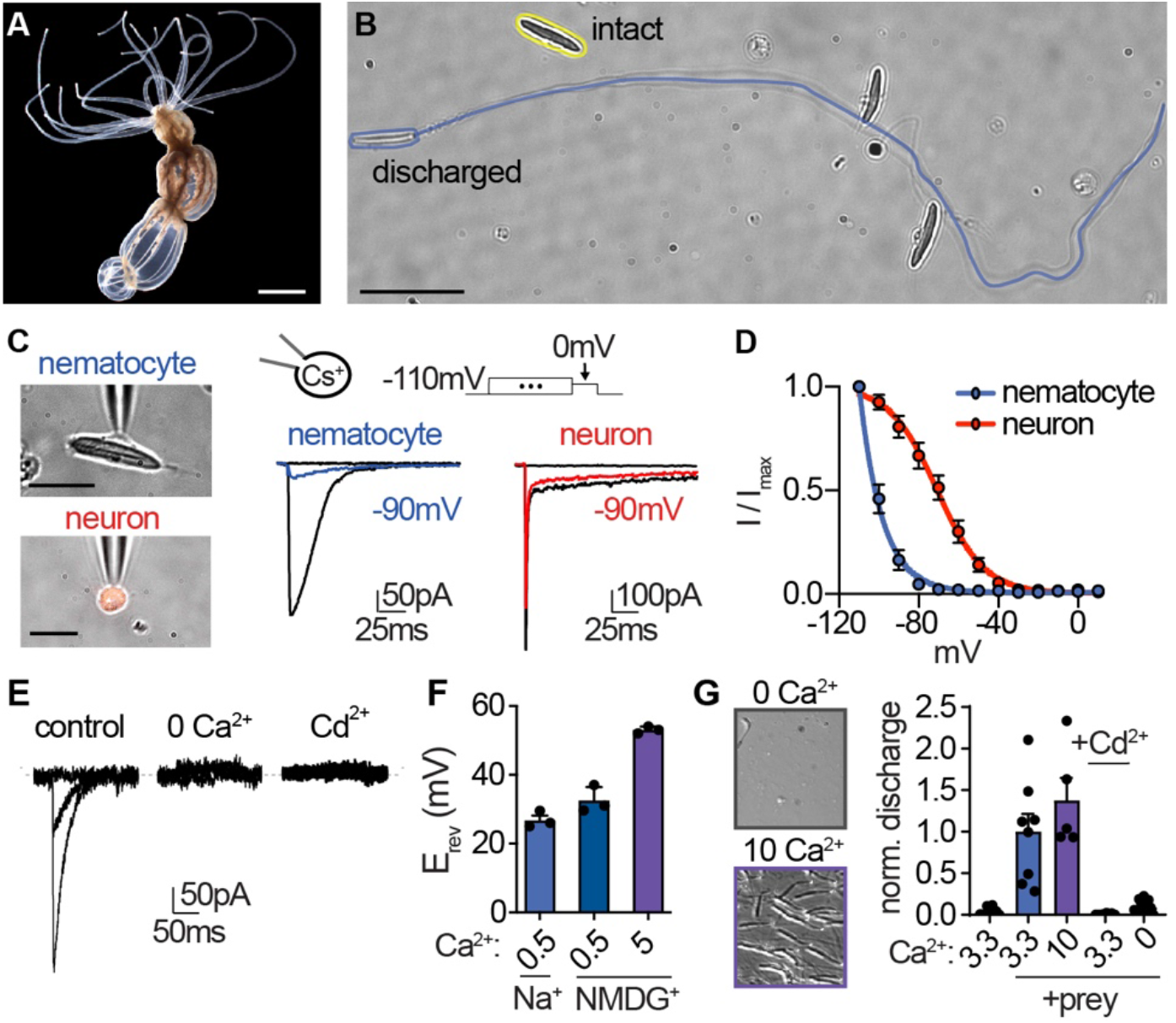
Nematocyte voltage-gated Ca^2+^ currents exhibit sensitive voltage-dependent inactivation. **(A)** Starlet sea anemone *(Nematostella vectensis).* Scale bar = 3mm. **(B)** Intact (yellow) and discharged nematocyte (blue). Scale bar = 20μm. **(C)** *Left:* Representative patch clamp experiments from a nematocyte and tentacle neuron. Scale bar = 10μm. *Right:* Nematocyte or neuron voltage-gated currents elicited by a maximally activating voltage pulse following 1s pre-pulses to −110mV (max current), −90mV (colored), or 0mV (inactivated, no current). **(D)** Nematocyte voltage-gated currents inactivated at very negative voltages compared with neurons. Nematocyte inactivation occurred at voltages more negative than could be measured compared with a sigmoidal inactivation relationship in neurons: nematocyte estimated V_i1/2_ = −100.2 ± 0.37mV, n = 13 and neuron V_i1/2_ = −70.81 ± 1.03mV, n = 9. Activation thresholds were similar (Fig. 1 - supplement 1A). **(E)** Nematocyte voltagegated currents elicited by −40mV and 0mV pulses were abolished in absence of external Ca^2+^ and blocked by cadmium (Cd^2+^). Representative of n = 4 for 0 Ca^2+^ and 3 for Cd^2+^, p < 0.001 paired two-tailed student’s t-test. **(F)** Nematocyte voltage-gated currents were Ca^2+^-sensitive as substitution of extracellular Ca^2+^ affected the reversal potential. n = 3 – 4, p <0.001 for 5mM Ca^2+^ versus other conditions, one-way ANOVA with post-hoc Tukey test. **(G)** Nematocyte discharge was minimal or absent in response to mechanical stimulation alone (n = 11, 3.3mM Ca^2+^). In the presence of prey extract, mechanically-evoked discharge was similar in standard and higher concentration of extracellular Ca^2+^ (n = 8 for 3.3mM Ca^2+^, n = 5 for 10mM) and blocked by Cd^2+^ (n = 8) or the removal of extracellular Ca^2+^ (n = 15). p <0.001 for + prey with 3.3 or 10mM Ca^2+^ versus other conditions, one-way ANOVA with post-hoc Bonferroni test. Data represented as mean ± sem.

Here, we demonstrate that nematocytes from *Nematostella vectensis* use a specialized Cav2.1 voltage-gated calcium channel orthologue (nCaV) to integrate dynamic voltage signals produced by distinct sensory stimuli. We show nematocytes are intrinsically mechanosensitive but nCa_v_ exhibits unique voltage-dependent inactivation that basally inhibits cellular activity, thereby preventing responses to extraneous mechanical stimuli, such as background water turbulence. We further show that sensory neurons make synaptic contact with nematocytes, and the neurotransmitter acetylcholine (ACh) elicits a hyperpolarizing response that relieves nCa_v_ inactivation to allow for subsequent cellular stimulation and chemo-tactile-elicited discharge. Thus, we propose that the specialized voltage dependence of nCa_v_ acts as a molecular filter for sensory discrimination.

## Results

### Nematocyte Ca_v_ channels

We first obtained whole-cell patch clamp recordings from acutely dissociated nematocytes using intracellular cesium (Cs^+^) to block potassium (K^+^) currents, thereby revealing a voltage-activated inward current (I_CaV_, **Fig. 1C**). In response to sustained positive voltage, voltage-gated ion channels typically enter a non-conducting, inactivated state and cannot be activated until returned to a resting state by negative membrane voltage. Remarkably, I_CaV_ began to inactivate at voltages more negative than we could technically measure, thus demonstrating an unusual voltage sensitivity of this conductance (**Fig. 1C, D**). To determine whether these properties were specific to nematocytes, we used a transgenic sea anemone with fluorescently-labeled neurons to facilitate direct comparison between these excitable cell types (Nakanishi et al., 2012) (**Fig. 1C**). Neuronal voltage-gated currents had a lower threshold for activation and exhibited much weaker voltage-dependent inactivation (**Fig. 1C, D, Fig. 1 - supplement 1A-D**), similar to currents found in neurons of other animals (Hille, 2001), indicating that nematocytes exhibit unusual voltage-dependent properties. Ion substitution and pore blocker experiments confirmed I_CaV_ is a Ca^2+^-sensitive current (**Fig. 1E, F**), consistent with the contribution of extracellular Ca^2+^ to chemo-tactile-induced discharge (Watson and Hessinger, 1994, Gitter et al., 1994) (**Fig. 1G**). Increased concentrations of extracellular Ca^2+^ did not affect inactivation (**Fig. 1 - supplement 1E**), suggesting the enhanced voltage-dependent inactivation is intrinsic to the channel complex. This observation is important because it suggests I_CaV_ renders nematocytes completely inactivated at typical resting membrane voltages and thus cells could not be stimulated from resting state.

To identify the ion channel mediating I_CaV_, we generated a tentacle-specific transcriptome and aligned reads from nematocyte-enriched cells (Sunagar et al., 2018). This strategy allowed us to search for differentially expressed transcripts that might encode Ca_v_ channel subunits (pore-forming a and auxiliary β and α2δ subunits). The orthologue of *cacnb2*, a β subunit of Ca_v_ channels, was the highest expressed Ca_v_ transcript in nematocyte-enriched cells, with levels 14-fold higher than other cells in the sea anemone (**Fig. 2A**). β subunits can modulate voltage-dependence and trafficking in diverse ways depending on their splice isoform, interacting subunits, and cellular context (Buraei and Yang, 2010). Importantly, β subunits only interact with α subunits of high voltage-activated (HVA) calcium channels (Perez-Reyes, 2003). In agreement with robust β subunit expression, we found significant enrichment for *cacna1a*, the pore-forming subunit of HVA CaV2.1, and high expression of *cacna2d1* (**Fig. 2 - supplement 1A, B**). These observations are consistent with a previous report demonstrating specific expression of *cacna1a* in nematocytes of sea anemone tentacles and expression of β subunits in nematocytes from jellyfish (Bouchard et al., 2006, Moran and Zakon, 2014, Bouchard and Anderson, 2014). Expression of *cacna1h*, which does not interact with auxiliary subunits (Buraei and Yang, 2010), was also observed, albeit at lower levels and across all cells (**Fig. 2 - supplement 1A**). Thus, it remains possible that voltage-gated currents in nematocytes are not carried exclusively by one Ca_v_ subtype.

**Figure 2:**
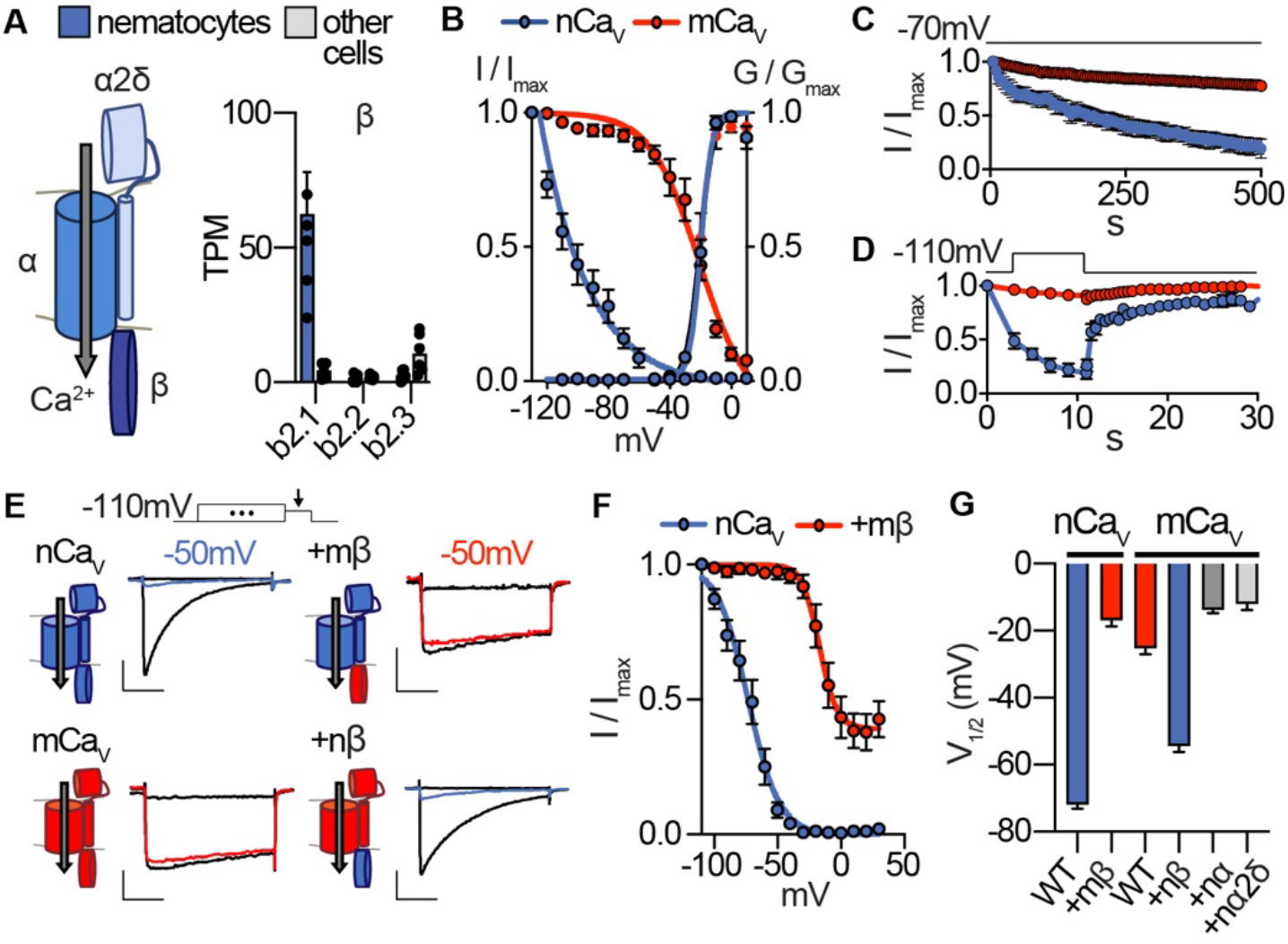
Nematostella Ca_v_ exhibits unique voltage-dependent properties conferred by its β subunit. **(A)** Ca_v_ channel complex with a, β, and α2δ subunits. mRNA expression (transcripts per million, TPM) of voltagegated calcium (Ca_v_) channel β subunits in nematocyte-enriched cells (blue) and non-enriched cells (grey). n = 6, p < 0.0001 for *cacnb2.1* in nematocytes versus other cells, two-way ANOVA with post-hoc Bonferroni test. **(B)** Heterologously-expressed nCav channels *(Nematostella cacna1a, cacnb2, cacna2d1)* inactivated at very negative voltages (estimated V_i1/2_ = −101.5 ± 1.60mV, n = 5) versus mammalian orthologues (mCa_v_, V_i1/2_ = −20.89 ± 3.35mV, n = 10). Activation thresholds were the same: nCa_v_ V_a1/2_ = −9.81 ± 0.28mV, n = 5, mCa_v_ V_a1/2_ = −10.41 ± 0.53mV, n = 9. Inactivation was measured in response to 1s pre-pulses from −110mV to 10mV with an inter-sweep holding potential of −90mV. **(C)** nCa_v_ exhibited slow inactivation with −70mV holding potential (0.2 Hz stimulation) that was best fit by two exponential functions with time constants of 9.97 and 369.5s. n = 6, multiple row two-tailed student’s t-test with significance of p < 0.05 by 15s and p < 0.0001 by 500s. **(D)** nCa_v_ inactivated at −40mV and quickly recovered at negative holding potentials. n = 7 for nCa_v_, n = 6 for mCa_v_. **(E)** Voltage-gated currents recorded from nCa_v_ or mCa_v_ following a −110mV pre-pulse, −50mV pre-pulse (colored), and 20mV pre-pulse. Ca_v_ β subunits were substituted as indicated (mammalian β in red and *Nematostella* β in blue). Scale bars = 100pA, 25ms. **(F)** Mammalian β shifts nCa_v_ voltage-dependent inactivation to positive voltages. nCa_v_ V_i1/2_ = −73.24 ± 1.17mV, n = 6. nCaV + mβ = −16.93 ± 1.85mV, n = 6. **(G)** Half maximal inactivation voltage (V_i1/2_) for Ca_v_ chimeras. p < 0.0001 for nCaV versus nCa_v_ + mβ, mCa_v_ versus mCa_v_ + nβ, one-way ANOVA with post-hoc Tukey test. Inactivation was measured in response to pre-pulses from −100mV to 10mV with an inter-sweep holding potential of −110mV to reduce slow inactivation. Data represented as mean ± sem.

Heterologous expression of *Nematostella* Ca_v_ (nCaV: *cacna1a, cacnb2*, and *cacna2d1)* produced voltage-gated currents with an activation threshold nearly identical to the Ca_v_ complex made from respective mammalian orthologues (mCa_v_, **Fig. 2B, Fig. 2 - supplement 1C, D**). Both channels had similar activation kinetics, but fast inactivation was significantly pronounced in nCa_v_, resembling native I_CaV_ (**Fig. 2E, Fig. 2 - supplement 1E**). Importantly, nCa_v_ voltage-dependent inactivation was greatly enhanced compared with mCa_v_, regardless of the charge carrier (**Fig. 2B, Fig. 2 - supplement 1F, G**). Similar to I_CaV_, nCa_v_ exhibited unusually-sensitive voltage-dependence and began to inactivate at voltages more negative than we could measure with an estimated midpoint inactivation voltage (V_i1/2_) ~80mV more negative than mCa_v_ (**Fig. 2B**). Even with a 5-second intersweep holding potential of −70mV, nCa_v_ exhibited slow inactivation resulting in a drastic decrease in channel availability over time (**Fig. 2C**). This slow inactivation was largely prevented by adjusting the holding potential to −110mV, suggesting inactivation occurs from the closed-state at potentials near or more negative than typical resting membrane potential (**Fig. 2 - supplement 1H**). Importantly, nCa_v_ rapidly recovered from inactivation, demonstrating that channels could be reset for subsequent activation following brief exposure to negative voltage (**Fig. 2D**). These distinctive features closely match the unique properties of native I_CaV_, suggesting nCa_v_ forms the predominant Ca_v_ channel in nematocytes.

To determine the molecular basis for nCa_v_ inactivation, we analyzed chimeric Ca_v_ complexes containing specific a, β, and α2δ1 subunits from nCa_v_ or mCa_v_ orthologues. Using an inter-sweep holding potential of −110mV to prevent slow inactivation, we compared voltage-dependent inactivation across chimeric channel complexes and found that only transfer of the β subunit significantly affected voltage-dependent inactivation, while a or a2δ1 subunits produced minimal effects on voltage-dependent activation, inactivation, or kinetics (**Fig. 2E, Fig. 2 - supplement 1C-E**). Indeed, other β subunits can induce significant hyperpolarized shifts in inactivation of HVA Ca_v_ channels (Yasuda et al., 2004). In this case, the mCa_v_ β subunit drastically shifted nCa_v_ inactivation by ~56mV in the positive direction, prevented complete inactivation, and produced slower fast inactivation (**Fig. 2E-G, Fig. 2 - supplement 1E**). Furthermore, nCa_v_ β was sufficient to confer greatly enhanced voltagedependent inactivation to mCa_v_ (**Fig. 2E, G**). From these results, we conclude that nCa_v_ β, the most enriched Ca_v_ subunit in nematocytes, confers nCa_v_s uniquely-sensitive voltage-dependent inactivation.

### Nematocyte excitability

Because electrical stimulation has been implicated in nematocyte discharge and some nematocytes can produce action potentials (Anderson and Mckay, 1987, Mckay and Anderson, 1988, Anderson and Bouchard, 2009), we used current-clamp to record the electrical responses of nematocytes to depolarizing stimuli. Under our conditions, nematocytes had a resting potential of −64.8 ± 8.9mV and did not produce a voltage spike when injected with current from rest (**Fig. 3A**). We further considered that the strong voltage-dependent inactivation of I_CaV_ could prevent excitability. Consistent with this idea, when nematocytes were first hyperpolarized to −90mV and subsequently stimulated by current injection, we observed a singular long voltage spike (**Fig. 3B, C**). In contrast, tentacle neurons produced multiple narrow spikes when injected with equivalent current amplitudes from a similar resting voltage, consistent with other neural systems (**Fig. 3A-C**). Differences in spike width and frequency appear suited to mediate distinctive cellular functions: dynamic information processing in neurons and a single robust discharge event in nematocytes. Furthermore, these results indicate strong voltage-dependent inactivation prevents nematocyte activation from rest.

**Figure 3:**
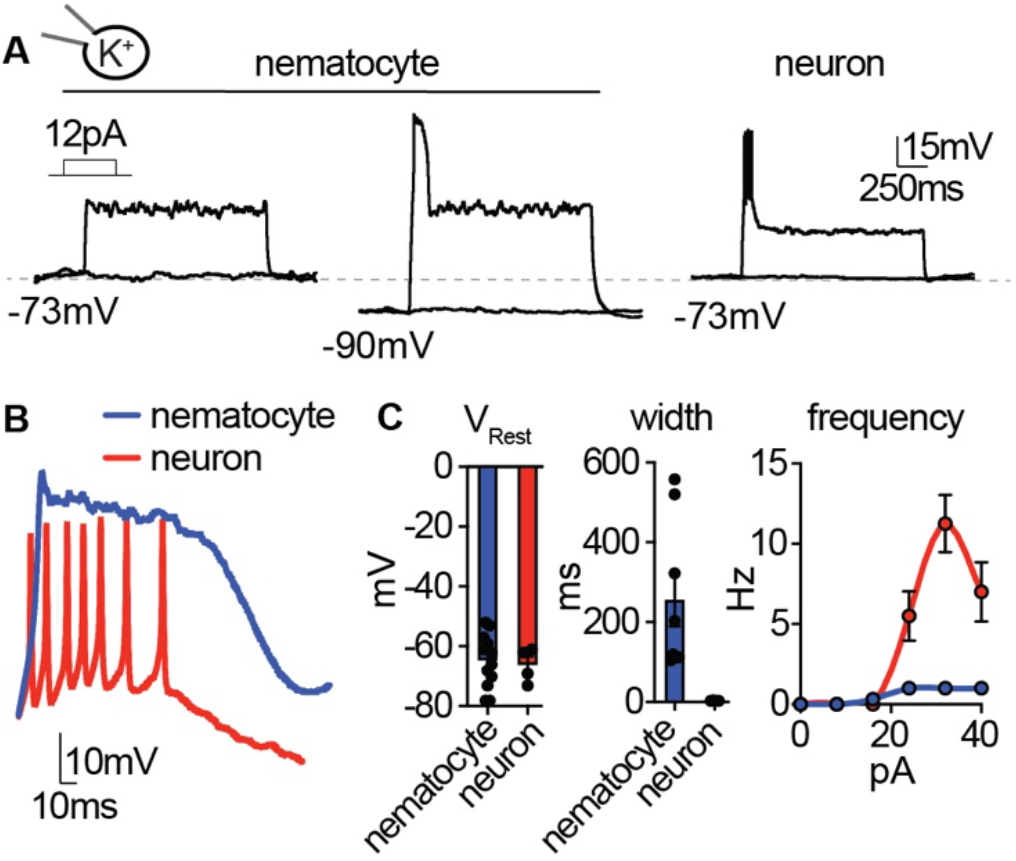
Nematocyte excitability requires hyperpolarized voltages. **(A)** Depolarizing current injection only elicited spikes from nematocytes first hyperpolarized to relieve inactivation. Nematocyte spike amplitude: 0mV at rest, 41.50 ± 2.20mV from ~-90mV, n = 8, p < 0.0001 two-tailed paired student’s t-test. In contrast, tentacle neurons spiked from rest (31.50 ± 1.71mV, n = 4). **(B)** Current injection elicited long singular spikes from nematocytes and numerous narrow spikes from neurons. **(C)** Nematocytes and neurons had similar resting membrane potentials but distinct spike width. n = 8 nematocytes, 4 neurons, p < 0.01, two-tailed student’s t-test. Nematocytes produced only one spike, regardless of injection amplitude (n = 8), whereas neurons produced varying spike frequency depending on injection amplitude (n = 4). p < 0.0001 two-way ANOVA with post-hoc Bonferroni test. Data represented as mean ± sem.

K^+^ channels often modulate repolarization following voltage spikes, so we compared K^+^ currents in nematocytes and tentacle neurons to understand how spike width might be differentially regulated. Nematocytes exhibited transient outward K^+^ currents that quickly inactivated, while neurons had large sustained K^+^ currents, perhaps important for repolarization and repetitive spiking (**Fig. 3 - supplement 1A-C**). The transient component of the nematocyte K^+^ current was highly sensitive to voltage-dependent inactivation, similar to I_CaV_ (**Fig. 3 - supplement 1D**). This K^+^ current was abolished by the rapid intracellular Ca^2+^ chelator BAPTA or by removing external Ca^2+^, similar to the effect of the K^+^ channel blocker TEA^+^ (**Fig. 3 - supplement 1E**). Consistent with this observation, nematocyte-enriched cells expressed numerous Ca^2+^-activated K^+^ channels (**Fig. 3 - supplement 1F**). Furthermore, intracellular Cs^+^ prolonged voltage spikes and greatly increased resting membrane voltage, substantiating a role for K^+^ currents in modulating membrane voltage (**Fig. 3 - supplement 1G**). We propose that these distinct K^+^ channel properties could contribute to the singular wide spikes of nematocytes versus the numerous narrow spikes of neurons.

### Nematocyte sensory transduction

If nematocytes are basally inhibited due to the unique voltage-dependent inactivation of I_CaV_, how do they respond to sensory signals to elicit discharge? Nematocyte discharge requires simultaneous detection of chemo- and mechanosensory cues (Pantin, 1942a, Watson and Hessinger, 1992), even though mechanical stimulation of the nematocyte’s cilium (cnidocil) within intact tentacles can by itself induce cellular depolarization (Brinkmann et al., 1996, Anderson and Bouchard, 2009). Indeed, we found the deflection of isolated nematocyte cnidocils elicited a mechanically-gated inward current with rapid activation and inactivation kinetics that was abolished by the mechanoreceptor blocker gadolinium (Gd^3+^,) and not observed in neurons (**Fig. 4A-C**). Furthermore, nematocyte-enriched cells differentially expressed transcripts encoding NompC (no mechanoreceptor potential C, **Fig. 4 - supplement 1A**), a widely conserved mechanoreceptor previously found to localize to the cnidocil of nematocytes from *Hydra* (Schuler et al., 2015). Heterologous expression of *Nematostella* NompC (nNompC) resulted in a mechanically-gated current with similar properties to native nematocytes, including rapid kinetics and Gd^3+^ sensitivity (**Fig. 4A-C**). Comparison with the *Drosophila* orthologue (dNompC) demonstrated nNompC had similar rapid kinetics, Gd^3+^ sensitivity, pressure-response relationships, and nonselective cation conductance, all consistent with the conservation of protein regions important for mechanosensitivity and ion selectivity (Jin et al., 2017) (**Fig. 4A-C, Fig. 4 - supplement 1B-D**). Thus, we conclude nematocytes are intrinsically mechanosensitive and suggest nNompC contributes to nematocyte mechanosensitivity. Importantly, this mechanically-evoked current is of sufficient amplitude to evoke a spike from very negative membrane voltages, but not from resting voltage at which I_CaV_ is inactivated.

**Figure 4:**
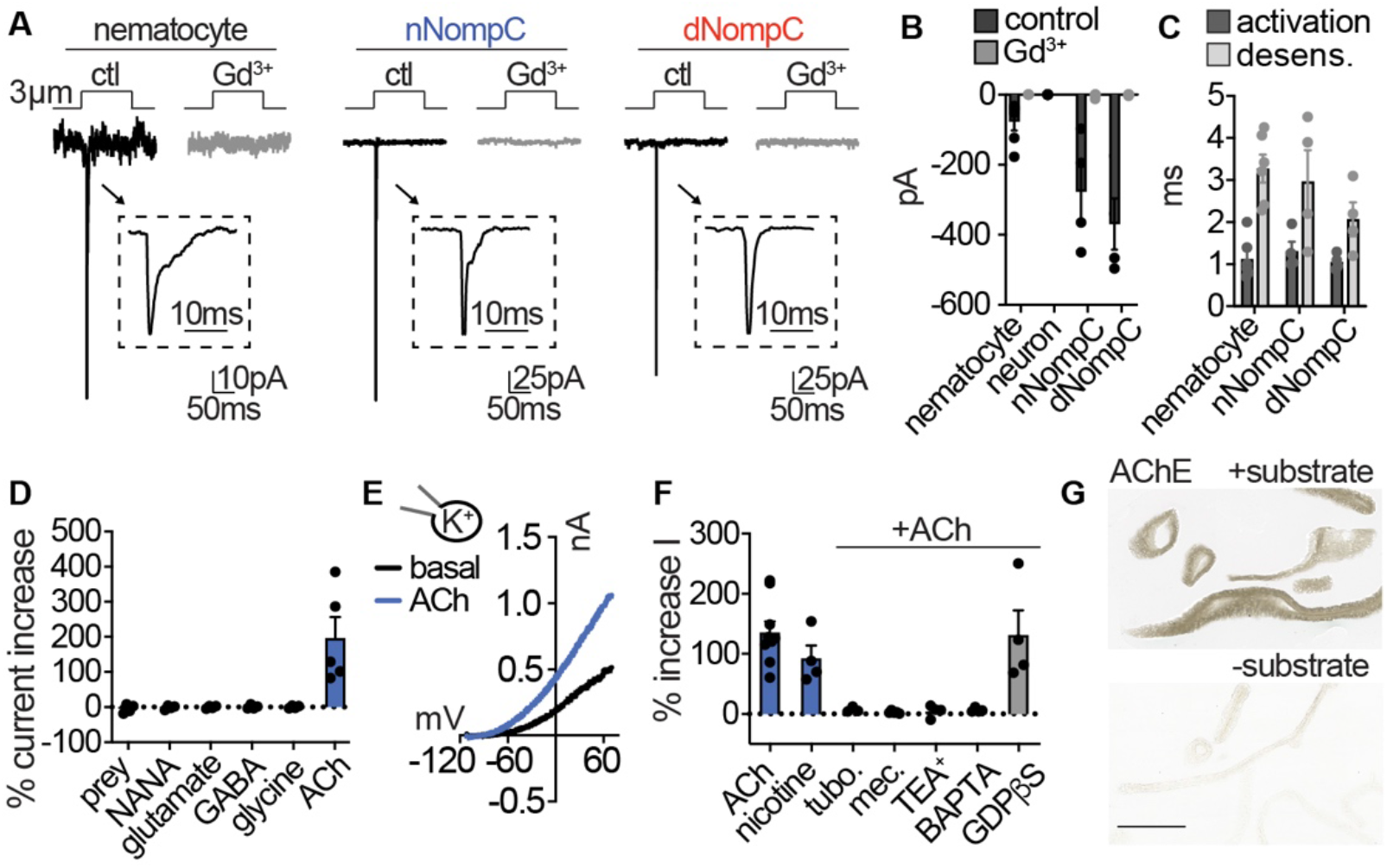
Nematocytes are intrinsically mechanosensitive and indirectly chemosensitive. **(A)** Mechanical stimulation of nematocytes evoked Gd^3+^-sensitive, transient inward currents with similar properties to heterologously-expressed *Nematostella* (n) and *Drosophila* (d) NompC channels. Stimulation thresholds (pipette displacement): nematocyte = 1.67 ± 0.33 μm (n = 6), nNompC = 4.23 ± 0.63 μm (n = 4), nNompC = 4.23 ± 0.48 μm (n = 4). **(B)** Mechanically-evoked currents from nematocytes (n = 6), nNompC (n = 4), and dNompC (n = 4) were blocked by Gd^3+^, while tentacle neurons lacked mechanically-evoked currents (n = 8). p < 0.01 two-tailed student’s t-test. **(C)** Mechanically-evoked current activation and desensitization kinetics were similar in nematocytes (n = 6), nNompC (n = 4), and dNompC (n = 4). **(D)** Chemosensory stimuli did not directly affect nematocytes but the neurotransmitter acetylcholine (ACh) elicited a large outward current. Prey extract = 4, NANA = 4, Glutamate = 4, GABA = 4, Glycine = 4, ACh = 5, p <0.001 for ACh versus other conditions, one-way ANOVA with post-hoc Tukey test. **(E)** Representative current-voltage relationship of ACh-elicited response in nematocytes. **(F)** ACh-evoked currents (n = 9) were blocked by nicotinic ACh receptor antagonists (tubocurarine = 4, mecamylamine = 4) and a similar current was elicited by nicotine (n = 4). ACh-evoked outward currents were inhibited by a K^+^ channel blocker (TEA^+^, n = 4) and an intracellular Ca^2+^ chelator (BAPTA, n = 4), but not the G-protein signaling blocker GDPβS (n = 4). p < 0.001 for vehicle versus antagonists, oneway ANOVA with post-hoc Tukey test. **(G)** Acetylcholinesterase staining in tentacles with and without substrate solution (representative of n = 3 animals). Scale bar = 200μm. Data represented as mean ± sem.

Because nematocyte discharge is mediated by combined chemical and mechanical cues (Pantin, 1942a, Watson and Hessinger, 1992), we wondered if chemosensory signals could modulate nematocyte membrane voltage to allow for I_CaV_ activation and cellular responses. While prey-derived chemicals evoked robust behavioral responses, similar treatments did not elicit electrical responses from isolated nematocytes (**Fig. 4D, Fig. 4 - supplement 1E**). Considering *in vivo* cellular and discharge activity requires the presence of prey-derived chemicals with simultaneous mechanical stimulation, chemoreception may occur indirectly through functionally-coupled cells (Price and Anderson, 2006). Previous studies suggest the presence of synaptic connections between nematocytes and other unknown cell types (Westfall et al., 1998, Oliver et al., 2008). To test this possibility, we screened isolated nematocytes for responses to well-conserved neurotransmitters and found only acetylcholine (ACh) evoked large outward currents (**Fig. 4D, E**). ACh-evoked currents were abolished by nicotinic acetylcholine receptor (nAChR) antagonists and recapitulated by nicotine (**Fig. 4F, Fig. 4 - supplement 1F**). The K^+^ channel blocker TEA^+^ and an intracellular Ca^2+^ chelator inhibited responses, but the G-protein signaling inhibitor GDPβS did not affect outward currents, consistent with the absence of muscarinic ACh receptors in *Nematostella* (Faltine-Gonzalez and Layden, 2019) (**Fig. 4F, Fig. 4 - supplement 1G**).

Furthermore, blockade of K^+^ currents with intracellular Cs^+^ revealed an ACh-elicited inward current was enhanced by increased extracellular Ca^2+^ and blocked by the nAChR antagonist mecamylamine (**Fig. 4 - supplement 1H**). In agreement with this observation, nematocyte- enriched cells expressed numerous nAChR-like transcripts which had well-conserved domains involved in Ca^2+^ permeability (Fucile, 2004) (**Fig. 4 - supplement 1I, J**). Finally, we found robust acetylcholinesterase activity in tentacles, further suggesting a role for ACh signaling in nematocyte function (**Fig. 4G**). These results demonstrate nematocytes use a nAChR-like signaling pathway to engage Ca^2+^-activated K^+^ channels, similar to how efferent cholinergic innervation of vertebrate hair cells modulates nAChR-K^+^ channel signaling to inhibit auditory responses (Elgoyhen and Katz, 2012).

To identify the origin of cellular connections to nematocytes, we used serial electron microscopy reconstruction to visualize nematocytes and neighboring cells. We analyzed similar tentacle tissues from which we carried out physiological experiments and readily observed neurons and nematocytes in close proximity (**Fig. 5 - supplement 1A, B**). In resulting micrographs, nematocytes were clearly identified by their distinct nematocyst and cnidocil (**Fig. 5 - supplement 1C**). Interestingly, each nematocyte exhibited a long process, of presently unknown function, that extended into the ectoderm (**Fig. 5 - supplement 1D**). We also observed numerous spirocytes, indicated by the presence of a large intracellular thread-like structure (**Fig. 5A, Fig. 5 - supplement 1C**). Putative sensory neurons were identified based on their synaptic contacts and extracellular projections (**Fig. 5A, Fig. 5 - supplement 1E, F**). Importantly, dense core vesicles were localized to electron-dense regions at the junction between each nematocyte and one other cell type, either sensory neurons or spirocytes (**Fig. 5A, Fig. 5 - supplement 2A-E**). Thus, nematocytes receive synaptic input from both neurons and spirocytes and likely serve as a site for integrating multiple signals (**Fig. 5 - supplement 2F, G**). This observation is consistent with the ability of cnidarians to simultaneously discharge multiple cnidocyte types to most efficiently capture prey (Pantin, 1942b).

**Figure 5:**
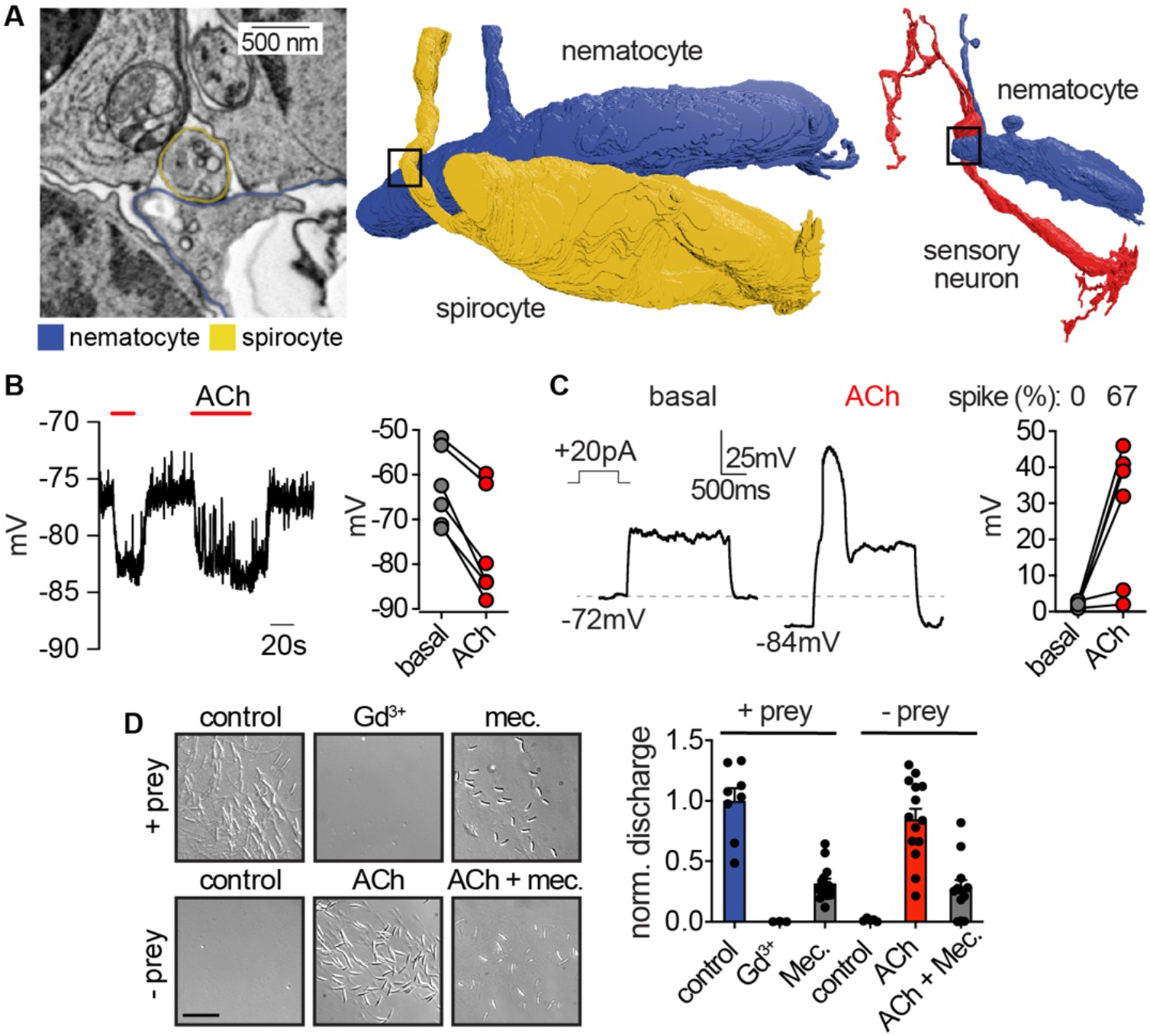
Nematocyte voltage-dependence facilitates signal integration required for stinging. **(A)** *Left:* Electron micrograph demonstrating dense-core vesicles in the vicinity of an electron-dense zone localized at the junction between a spirocyte and nematocyte. *Middle:* 3D reconstruction of the nematocyte and spirocyte shown in the left panel. The box indicates the position of the synapse. *Right:* 3D reconstruction of a different nematocyte making a synapse with a putative sensory neuron. The box indicates the position of the synapse shown in Fig. 5 - supplement 2C and E. **(B)** ACh induced hyperpolarization of nematocytes. n = 6, p < 0.01 paired two-tailed student’s t-test. **(C)** Depolarizing current injection did not induce active properties from rest, but did elicit a voltage spike from most nematocytes hyperpolarized by ACh. n = 6, p < 0.01 paired two-tailed student’s t-test. **(D)** Chemo-tactile-induced nematocyte discharge (touch + prey extract, n = 8) was inhibited by the mechanoreceptor blocker Gd^3+^ (n = 7) and nAChR antagonist mecamylamine (n = 14). In the absence of chemical stimulation (touch - prey extract, n = 5), touch + ACh (n = 14) was sufficient to induce discharge, which was inhibited by mecamylamine (n = 12). p < 0.0001 for controls versus respective treatments, one-way ANOVA with post-hoc Bonferroni test. Data represented as mean ± sem.

*Voltage-dependence mediates signal integration* How do distinct mechanosensory and chemosensory signals converge to elicit discharge? In agreement with our observation that nAChR activation increases K^+^ channel activity, ACh hyperpolarized nematocytes to negative voltages from which they were capable of producing robust voltage spikes (**Fig. 5B, C**). A select number of cells with a more positive resting voltage still failed to produce spikes, and therefore additional regulation could exist through the modulation of resting membrane voltage (**Fig. 5 - supplement 3A, B**). These results suggest that the voltage-dependence for I_CaV_ prevents basal activation to depolarizing signals, such as mechanical stimulation, but activation of nAChR hyperpolarizes the cell to relieve I_CaV_ inactivation, thereby amplifying depolarizing signals to mediate cellular responses.

Consistent with a requirement for both mechano- and chemosensory input, we found the mechanoreceptor blocker Gd^3+^ inhibited chemo-tactile stimulation of discharge. Additionally, the nAChR antagonist mecamylamine greatly reduced chemo-tactile-induced discharge (**Fig. 5D**). Washout of both treatments recovered the ability of nematocytes to discharge (**Fig. 5 - supplement 3C**). Moreover, the requirement for prey-derived chemicals was completely recapitulated by ACh (**Fig. 5D**). These results are consistent with a role for ACh signaling downstream of chemosensory stimulation. Thus, we propose that the unique I_CaV_ voltage-dependent inactivation provides a mechanism by which nematocytes filter extraneous depolarizing mechanical signals, but can integrate chemosensory-induced hyperpolarization together with a depolarizing stimulus to elicit robust signal amplification and discharge responses (**Fig. 6**).

**Figure 6:**
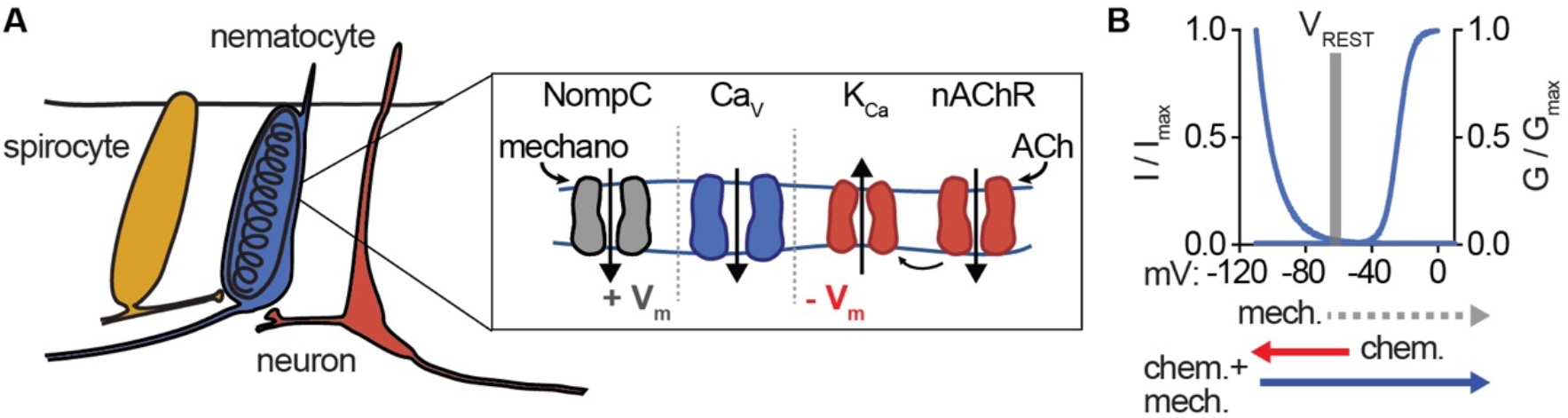
Nematocyte Ca_v_ properties filter salient chemo-tactile signals. **(A)** Model for nematocyte signal integration. **(B)** Nematocyte Ca_v_ is inactivated at rest and thus does not amplify extraneous NompC-mediated mechanical signals. Chemosensory stimuli hyperpolarize nematocytes through ACh signaling to relieve inactivation of Ca_v_ channels, which can then amplify mechanical stimuli to engage discharge.

## Discussion

Here, we demonstrate that nematocytes use the specialized properties of nCa_v_ to filter salient chemo-tactile signals from environmental noise. The involvement of Ca^2+^ signaling in this process is consistent with a well-established role for Ca^2+^ influx in mediating discharge across multiple cnidarian species (Mckay and Anderson, 1988, Santoro and Salleo, 1991, Watson and Hessinger, 1994). However, the exact mechanism by which Ca^2+^ influx mediates discharge is unclear. It has been proposed that Ca^2+^ influx alters the permeability of the nematocyst capsule and/or initiates the rapid dissociation of ions within the cyst to induce osmotic changes within the organelle (Lubbock and Amos, 1981, Lubbock et al., 1981, Weber, 1990, Tardent, 1995). Our transcriptomic analyses demonstrate that nematocytes express multiple Ca^2+^ handling proteins (**Fig. 5 - supplement 3D-F**), which could control Ca^2+^ signaling domains in response to spatially-restricted sensory transduction cascades. This organization would be consistent with the necessity for I_CaV_-mediated amplification of receptor-mediated nonselective cation conductances. Indeed, Ca_v_ currents could mediate an increase of >500μM Ca^2+^ considering a uniform distribution across the small cytoplasmic volume (5%) not occupied by the nematocyst. Future studies will provide insight into the coupling between sensory transduction and organellar physiology.

Our results demonstrate one mechanism by which nematocytes integrate combined mechanical and chemical cues to filter salient environmental information and appropriately engage nematocyte discharge. However, cnidarians occupy distinct ecological niches and may have evolved different biophysical features to account for increased turbulence, specific behaviors, or particular prey and predatory targets. Numerous sea anemones occupy turbulent tidal pools, whereas others, like *Nematostella vectensis*, live in calmer regions. Similarly, cnidarians can undergo developmental transitions between immobile and free-floating medusa phases while maintaining the use of nematocytes for prey capture (Martin and Archer, 1997). Although forces generated from swimming prey are likely negligible in comparison with physical contact of the cnidocil, strong tidal waves may be sufficient to elicit mechanoreceptive responses which could interfere with pertinent chemo-tactile sensation and subsequent stinging responses. These ecological differences might require distinct filtering mechanisms for distinguishing salient prey or predator signals. For example, anatomical organization and pharmacological dependence for cellular and discharge activity in anthozoan nematocytes differs from hydrozoans (Anderson and Mckay, 1987, Kass-Simon and Scappaticci, 2002, Oliver et al., 2008). Indeed, nematocysts vary extensively in morphology, differing in the length of the extruded thread, the presence of spines, and the composition of toxins (Kass-Simon and Scappaticci, 2002), likely reflecting the diversity of organismal needs.

The modularity provided by synaptic connections could increase the diversity of signals which modulate nematocyte discharge. For instance, distinct chemoreceptor cells could regulate specific nematocyte populations as some species of sea anemones possess specialized tentacles for attacking other anemones, while a separate set of tentacles are used for stinging prey (Brace, 1990). Discharge can be also regulated by organismal nutritional state (Sandberg et al., 1971), suggesting nematocytes could receive input from digestive cells or hormones. Additionally, various cnidocyte types and uses are found across the cnidarian phylum. Within anthozoans, nematocytes and spirocytes may use similar or distinct mechanisms to control discharge. Comparatively, the freshwater cnidarian, *Hydra vulgaris*, uses a specific cnidocyte to grasp surfaces for phototaxis, suggesting that these cells could be regulated downstream of a photoreceptor (Plachetzki et al., 2012). Functional comparisons will reveal whether specific proteins, domains, or signaling mechanisms are conserved or give rise to the evolutionary novelties across these incredibly specialized cell types (Babonis and Martindale, 2014).

The ability to distinguish behaviorally-relevant stimuli, such as prey, from background noise is especially critical because nematocytes are singleuse cells that must be replaced following discharge. Multiple species have taken advantage of these specialized conditions by adapting to evade and exploit nematocyte discharge for their own defensive purposes. For example, clownfish can live among the tentacles of sea anemones without harmful effects, although the exact mechanism by which this occurs is unclear (Lubbock, 1980). Certain species of nudibranchs and ctenophores acquire undischarged nematocysts from prey and store them for later defense, indicating that these organisms are able to initially prevent discharge responses (Greenwood, 2009). Understanding such regulation could reveal additional mechanisms by which cells process diverse stimuli and provide insight into the evolution of these interspecies relationships.

## Materials and Methods

### Animals and Cells

Starlet sea anemones *(Nematostella vectensis)* were provided by the Marine Biological Laboratory (Woods Hole, Massachusetts), Nv-Elav1::mOrange transgenic animals were a gift from F. Rentzsch. Adult animals of both sexes used for cellular physiology experiments were fed freshly hatched artemia twice a week and kept on a 12 hr light/dark cycle in 1/3 natural sea water (NSW). Nematocytes and neurons were isolated from tentacle tissue, which was harvested by anesthetizing animals in high magnesium solution containing (mM): 140 NaCl, 3.3 Glucose, 3.3 KCl, 3.3 HEPES, 40 MgCl_2_. Cells were isolated from tentacles immediately prior to electrophysiology experiments by treatment with 0.05% Trypsin at 32°C for 30 min and mechanical dissociation in divalent free recording solution (mM): 140 NaCl, 3.3 Glucose, 3.3 KCl, 3.3 HEPES, pH 7.6. Basitrichous isorhiza nematocytes were isolated from tentacles and identified by a thick-walled capsule containing a barbed thread, with a characteristic high refractive index, oblong shape and presence of a cnidocil. Spirocytes were identified by a thin-walled capsule containing a thin, unarmed thread, used for ensnaring prey. Neurons were identified by mOrange expression.

HEK293T cells (ATCC) were grown in DMEM, 10% fetal calf serum, and 1% penicillin/streptomycin at 37°C, 5% CO_2_. Cells were transfected using lipofectamine 2000 (Invitrogen/Life Technologies) according to the manufacturer’s protocol. 1 μg of *Nematostella cacna1b1, cacnb2.1, cacna2d1.1. M. musculus* (mouse) *cacna1a, R. norvegicus* (rat) *cacnb2a*, or rat *cacna2d1* was coexpressed with 0.5 μg GFP. Mechanosensitive proteins were assayed using HEK293T cells transfected with 1 μg of either *Drosophila* NompC or *Nematostella* NompC. To enhance channel expression, cells were transfected for 6-8h, plated on coverslips, and then incubated at 28°C for 2-6 days before experiments. Rat *cacna2d1* and *cacna1a* were gifts from D. Lipscombe (Addgene plasmids 26575 and 26578) and *cacnb2a* was a gift from A. Dolphin (Addgene plasmid 107424). *Drosophila* NompC-GFP was a gift from YN Jan.

### Molecular biology

RNA was prepared from tentacles of adult *Nematostella* using published methods (Stefanik et al., 2013). Briefly, 50mg of tentacle tissue were homogenized and RNA was extracted using TRIzol. RNA was isolated and DNase treated (Zymo Research), then used for cDNA library synthesis (NEB E6560). Full-length sequence for a *Nematostella* calcium channel beta subunit was obtained with a RACE strategy using specific primers (GATTACGCCAAGCTTTATGCGTCCAATCGTACTTGTCGGC and GATTACGCCAAGCTTGCCGACAAGTACGATTGGACGCATA) on the amplified tentacle-tissue library. The final sequence was confirmed using primers corresponding to the end of the derived sequence (CAGAGCCAGGCCTGAGCGAG and GCCCCGTTAAAAGTCGAGAG) to amplify a fulllength cDNA from tentacle mRNA, which was sequenced to confirm identity. *Cacna1a, cacna2d1, cacnb2*, and nNompC-GFP were synthesized by Genscript (Piscataway, NJ). Sequence alignments were carried out using Clustal Omega.

### Transcriptomics

Tentacle tissue was ground to a fine powder in the presence of liquid nitrogen in lysis buffer (50 mM Tris-HCl pH 7.5, 250 mM KCl, 35 mM MgCl_2_, 25 mM EGTA-KOH pH 8, 5 mM DTT, murine RNase inhibitor (NEB), 1% (w/v) NP-40, 5% (w/v) sucrose, 100 μg ml^−1^ cycloheximide (Sigma), 500 μg ml^−1^ heparin (Sigma)). Lysate was incubated on ice for 5 min, triturated 5 times with an 18 g needle, and insoluble material was removed by centrifugation at 16000g for 5 min at 4 °C. Polyadenylated RNA was used to make sequencing libraries and sequenced on an Illumina HiSeq 4000 (Novogene). Quality filtering and adapter trimming was performed using Cutadapt (Martin, 2011), and a *de novo* transcriptome was assembled using Trinity (Grabherr et al., 2011). Annotation was performed using InterProScan (Jones et al., 2014) with Panther member database analysis, HMMER (Eddy, 2009) with the Pfam (El-Gebali et al., 2019) database, and DIAMOND (Buchfink et al., 2015) with the UniProt/TrEMBL database.

Reads from sorted cnidocytes (Bioproject PRJNA391807 (Sunagar et al., 2018)), enriched for nematocytes, were quality and adapter trimmed as described above, and transcript abundance (TPM) was quantified using Kallisto (Bray et al., 2016) and the tentacle transcriptome. For read mapping visualization, mapping was performed with Bowtie2 (Langmead and Salzberg, 2012), output files were converted to indexed bam files using Samtools (Li et al., 2009), and visualization was performed with the Integrated Genomics Viewer (Robinson et al., 2011).

### Electrophysiology

Recordings were carried out at room temperature using a MulticClamp 700B amplifier (Axon Instruments) and digitized using a Digidata 1550B (Axon Instruments) interface and pClamp software (Axon Instruments). Whole-cell recording data were filtered at 1kHz and sampled at 10kHz. For single-channel recordings, data were filtered at 2kHz and sampled at 20kHz. Data were leak subtracted online using a p/4 protocol, and membrane potentials were corrected for liquid junction potentials. For whole-cell nematocyte and neuron recordings, borosilicate glass pipettes were polished to 8-10MΩ. The standard *Nematostella* medium was used as the extracellular solution and contained (in mM): 140 NaCl, 3.3 glucose, 3.3 KCl, 3.3 HEPES, 0.5 CaCl_2_, 0.5 MgCl_2_, pH 7.6. Two intracellular solutions were used for recording. For isolating inward currents (mM): 133.3 cesium methanesulfonate, 1.33 MgCl_2_, 3.33 EGTA, 3.33 HEPES, 10 sucrose, 10 CsEGTA, pH 7.6. For outward currents (mM) 166.67 potassium gluconate, 3.33 HEPES, 10 sucrose, 1.33 MgCl_2_, 10 KEGTA, pH 7.6. In some experiments, BAPTA was substituted for EGTA. For whole-cell recordings in HEK293 cells, pipettes were 3-4MΩ. The standard extracellular solution contained (in mM): 140 NaCl, 5 KCl, 10 HEPES, 2 CaCl_2_, 1 MgCl_2_, pH 7.4. The intracellular solution contained (mM): 140 cesium methanesulfonate, 1 MgCl_2_, 3.33 EGTA, 3.33 HEPES, 10 sucrose, pH 7.2. In ion substitution experiments, equimolar Ba^2+^ was substituted for Ca^2+^. Single-channel recording extracellular solution contained (mM): 140 NaCl, 10 HEPES, 1 NaEGTA, pH 7.4. The intracellular solution used (mM): 140 CsCl, 10 HEPES, 1 CsEGTA, pH 7.4.

The following pharmacological agents were used: N-Acetylneuraminic acid (NANA, 100μM, Sigma), glycine (100mM, Sigma), acetylcholine (1mM), mecamylamine (100μM, 500μM for behavioral experiments, Tocris), GdCl_3_ (100μM, Sigma), glutamate (1mM), GABA (1mM), nicotine (100μM, Tocris), tubocurarine (10μM, Tocris), TEA (10mM, Sigma), BAPTA (10mM, Tocris), GDPβS (1mM, Sigma), and Cd^2+^ (500μM, 250 μM for behavioral experiments). All were dissolved in water. Pharmacological effects were quantified as differences in normalized peak current from the same cell following bath application of the drug (I_treatment_/I_control_). Whole-cell recordings were used to assess mechanical sensation together with a piezoelectric-driven (Physik Instrumente) fire-polished glass pipette (tip diameter 1μm). Mechanical steps in 0.5μm increment were applied every 5s while cells were voltage-clamped at −90mV. Single mechanosensitive channels were studied using excised outside-out patches exposed to pressure applied via a High-Speed Pressure Clamp system (HSPC, ALA-scientific). Pressure-response relationships were established using pressure steps in 10mmHg increments. Voltage-dependence of currents was measured from −100mV to 100mV in 20mV increments while applying repetitive 60mmHg pressure pulses.

Unless stated otherwise, voltage-gated currents were measured in response to a 200ms voltage pulse in 10mV increments from a −110mV holding potential. G-V relationships were derived from I-V curves by calculating G: G=I_CaV_/(V_m_-E_rev_) and fit with a Boltzman equation. Voltage-dependent inactivation was measured during −10mV (Ca^2+^ currents in native cells), 0mV (Ca^2+^ currents in heterologously expressed channels), 60mV (K^+^ currents in native cells) voltage pulses following a series of 1s pre-pulses ranging from −110mV to 60mV. Voltage-dependent inactivation was quantified as I/I_max_, with I_max_ occurring at the voltage pulse following a −110mV prepulse. In some instances, inactivation curves could not be fit with a Boltzman equation and were instead fitted with an exponential. The time course of voltage-dependent inactivation was measured by using a holding voltage of −110mV or −70mV and applying a 0mV test pulse every 5s. Recovery from inactivation was quantified by normalizing inactivated and recovered currents to those elicited from the same cell in which a 0mV voltage pulse was applied from −110mV. Test pulses from a holding voltage of −40mV were used to assess inactivation, followed by pulses from −110mV where currents quickly recovered to maximal amplitude. Repetitive stimulation using 20ms pulses to −10mV or 0mV from a holding voltage of −90mV was also used to measure inactivation in response to repetitive stimulation. Current inactivation kinetics were quantified by the portion of current remaining at the end of 200ms pulse (R200) or fit with a single exponential. Activation was quantified as the time from current activation until peak amplitude. 200ms voltage ramps from – 120 to 100mV were used to measure ACh-elicited currents. Stimulus-evoked currents were normalized to basal currents measured at the same voltage of 80mV.

Single channel currents were measured from the middle of the noise band between closed and open states or derived from all-points amplitude histograms fit with Gaussian relationships at closed and open peaks for each excised patch record. Conductance was calculated from the linear slope of I-V relationships. N(P_o_) was calculated during pressure steps while voltage was held at −80mV. In current clamp recordings, effects of ACh or intracellular ions on resting membrane potential was measured without current injection (I=0). 1s depolarizing current steps of various amplitudes were injected to measure spikes which were quantified by frequency (spikes/second) or width (duration of spike). To test whether resting membrane potential affects the ability to generate spikes, hyperpolarizing current was injected to bring cells to negative voltages (<-90mV) or ACh was locally perfused before depolarizing current injection.

The change in Ca^2+^ concentration from a nematocyte voltage spike was estimated based on the integral of Ca^2+^-selective nematocyte currents elicited by a 0mV step, the same amplitude and slightly shorter duration than a voltage spike. Nematocyte volume was estimated from serial electron microscopy reconstruction with a non-nematocyst volume of approximately 5% of the total volume of the cell. We did not consider the volume occupied by other organelles, making for a conservative estimate. Furthermore, calculations were made with extracellular recording solution containing 0.5mM Ca^2+^, which is approximately 6-fold less than physiological concentrations. Thus, the large increase we calculated likely underestimates the total Ca^2+^ influx.

### Immunohistochemistry

Neural staining: Adult Nv-Elav1::mOrange *Nematostella* were paralyzed in anesthetic solution, then placed in a 4% solution of PFA overnight. Animals were cryoprotected using a gradient of increasing sucrose concentrations (10% to 50%) in PBS over two days. Cryostat sections (20μm thick) were permeabilized with 0.2% Triton-X and 4% normal goat serum (NGS) at room temperature for 1 hour, followed by incubation with DsRed Polyclonal Antibody (Takara Bio # 632496) overnight in PBST (0.2%) and NGS at 4°C. Tissue was rinsed 3 times with PBST before secondary was applied (Goat antirabbit 647, Abcam) for 2 hours at room temperature. Tissue was rinsed with PBS and mounted with Vectashield containing DAPI (Novus Biologicals).

Acetylcholinesterase staining: Tentacles were stained for the presence of acetylcholinesterase as described (Cold Spring Harbor protocol: Staining for Acetylcholinesterase in Brain Sections) using 40μm thick cryosections mounted on glass slides. Slides were incubated in acetylthiocholine and copper-buffered solution at 40°C until tentacles appeared white. The stain was developed with a silver solution so that stained areas appear brown. Slides were incubated in the presence of the silver staining solution (+substrate) or saline (-substrate), rinsed according to protocol, and mounted in Fluoromount-G (SouthernBiotech) and imaged using a scanning, transmitted light microscope.

### Behavior

Discharge of nematocysts was assessed based on well-established assays (Watson and Hessinger, 1994, Gitter et al., 1994). Adult *Nematostella* were placed in petri dishes containing a modified *Nematostella* medium, containing 16.6mM MgCl_2_. Animals were given appropriate time to acclimate before presented with stimuli. For assaying discharge, 5 mm round coverslips were coated with a solution of 25% gelatin (w/v) dissolved in medium, and allowed to cure overnight prior to use. Coverslips were presented to the animal’s tentacles for 5 seconds and then immediately imaged at 20X magnification using a transmitted light source. To assay behavioral response to prey-derived chemicals, freshly hatched brine shrimp were flash frozen and pulverized, then filtered through a 0.22μm filter. Coverslips were dipped in the prey extract and immediately presented to the animal. All pharmacological agents were bath applied, except for acetylcholine (1mM), which was delivered as a bolus immediately prior to coverslip presentation. Acetylcholine exposure did not produce movement or contraction of tentacles. Experiments carried out in the absence of extracellular Ca^2+^ were nominally Ca^2+^ free and did not use extracellular chelators. The highest density of discharged nematocytes on the coverslip was imaged at 20X. Images were blindly analyzed using a custom Matlab routine (available in supplemental material) in which images were thresholded and the fraction of pixels corresponding to nematocytes was compared across experiments.

### Electron microscopy

Tentacles from an individual *Nematostella vectensis* were placed between two sapphire coverslips separated by a 100um spacer ring (Leica) and frozen in a high-pressure freezer (EM ICE, Leica). This was followed by freeze-substitution (EM AFS2, Leica) in dry acetone containing 1 % ddH2O, 1 % OsO4 and 1 % glutaraldehyde at −90°C for 48 hrs. The temperature was then increased at 5°C/h up to 20°C and samples were washed at room temperature in pure acetone 3×10min RT and propylene oxide 1×10min. Samples were infiltrated with 1:1 Epon: propylene oxide overnight at 4C. The samples were subsequently embedded in TAAB Epon (Marivac Canada Inc.) and polymerized at 60 degrees C for 48 hrs. Ultrathin sections (about 50nm) were cut on an ultramicrotome (Leica EM UC6) and collected with an automated tape collector (ATUM (Kasthuri et al., 2015)). The sections were then post-stained with uranyl acetate and lead citrate prior to imaging with a scanning electron microscope (Zeiss SIGMA) using a back-scattered electron detector and a pixel size of 4nm.

Once all the sections were scanned, images were aligned into a stack using the algorithm “Linear Stack Alignment with SIFT” available in Fiji (Schindelin et al., 2012). After alignment, images were imported into VAST (Berger et al., 2018) so that every cell could be manually traced. By examining sections and following cellular processes contacts between cells of interest (e.g. neurons and nematocytes) were identified and assessed for the presence of dense-core vesicles in the vicinity (~500nm). Such instances were labeled as putative synapses. Cells were then rendered in 3 dimensions using 3D Studio Max 2019 (Autodesk, San Rafael, CA).

Nematocytes were readily identified because resin does not infiltrate the nematocyst capsule, making for an ‘empty’ appearance (large white area). Spirocytes were also readily identified based on their capsule containing a long, coiled filament. Sensory neurons were identified according to the higher number of dense core vesicles, higher number of synapses and the presence of sensory processes extending into the external environment.

### Statistical analysis

Data were analyzed with Clampfit (Axon Instruments) or Prism (Graphpad) and are represented as mean ± s.e.m. *n* represents independent experiments for the number of cells/patches or behavioral trials. Data were considered significant if p < 0.05 using paired or unpaired two-tailed Student’s t-tests or one- or twoway ANOVAs. All significance tests were justified considering the experimental design and we assumed normal distribution and variance, as is common for similar experiments. Sample sizes were chosen based on the number of independent experiments required for statistical significance and technical feasibility.

### Data availability

Deep sequencing data will be archived under Gene Expression Omnibus and GenBank accession numbers will be submitted upon publication. All plasmids are available upon request. Further requests for resources and reagents should be directed to and will be fulfilled by the corresponding author, NWB.

## Acknowledgments

We thank J. Lichtman and F. Engert for helpful suggestions throughout this study, B. Bean, D. Julius, and R. Nicoll for critical reading of the manuscript, K. Boit for assistance with electron microscopy analyses, J. Turecek for assistance with immunohistochemistry, and K. Koenig for help with sea anemones. This research was supported by grants to NWB from the New York Stem Cell Foundation, Searle Scholars Program, Alfred P. Sloan Foundation, Klingenstein-Simons Fellowship, and the NIH (R00DK115879), as well as the Swiss National Science Foundation (P2SKP3_187684 / 1) to CD and NIH (U24NS109102) to J. Lichtman.

## Author Contributions

KW, LVG, ASYL, and NWB contributed to molecular and physiological studies. CP contributed to anatomical studies. All authors were involved with writing or reviewing the manuscript.

## Competing Interests

The authors declare no competing financial interests.

---

**Figure 1 - figure supplement 1:**
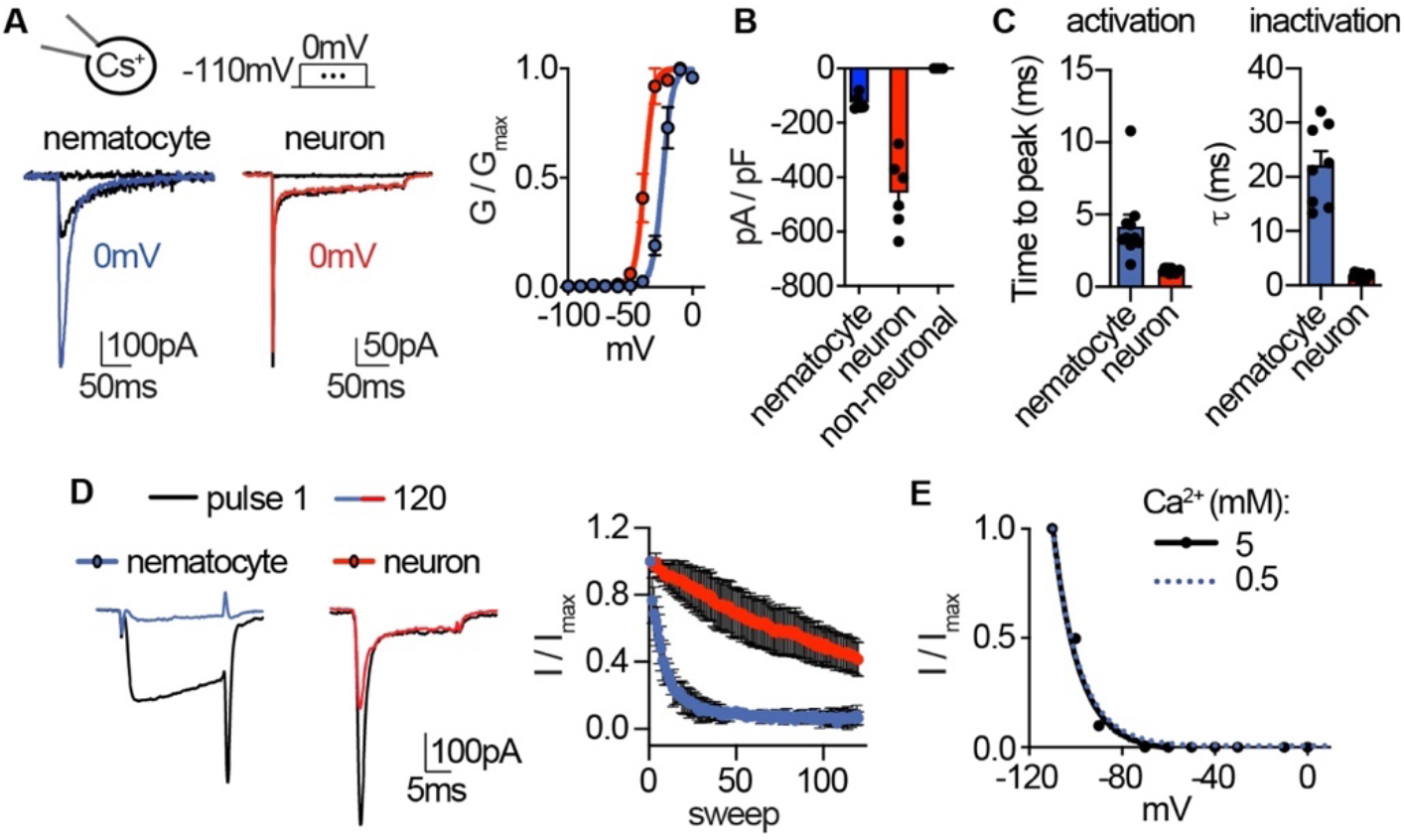
Native nematocyte Ca_v_ properties. **(A)**Voltage-gated currents from a nematocyte and tentacle neuron elicited by −40mV or 0mV pulses. Conductance-voltage curves: nematocyte V_a1/2_ = −24.02 ± 0.53mV, n = 10 and neuron V_a1/2_ = −38.59 ± 0.64mV, n = 7. **(B)** Peak current density elicited by 0mV pulse in nematocytes (n = 5), neurons (n = 6), and non-neuronal cells (n = 3). **(C)** Inward currents activated and inactivated more slowly in nematocytes compared with neurons. n = 10 nematocytes, 11 neurons. p < 0.0001 two-tailed student’s t-test. **(D)** Nematocyte inward currents inactivated more quickly with repetitive stimulation compared with neurons. Protocol: 3.33 Hz stimulation with 20ms pulses to −10mV from −90mV. n = 4 of each cell type, multiple row two-tailed student’s t-test with significance of p < 0.05 by sweep 5 and p < 0.0001 by 120. **(E)** Inactivation of inward currents does not vary with external Ca^2+^. 0.5 mM Ca^2+^ estimated V_i1/2_ = −100.2 ± 0.36mV, n = 13, from Fig 1. 5 mM Ca^2+^ estimated V_i1/2_ = −99.79 ± 0.87mV, n = 3. Data represented as mean ± sem.

**Figure 2 - figure supplement 1:**
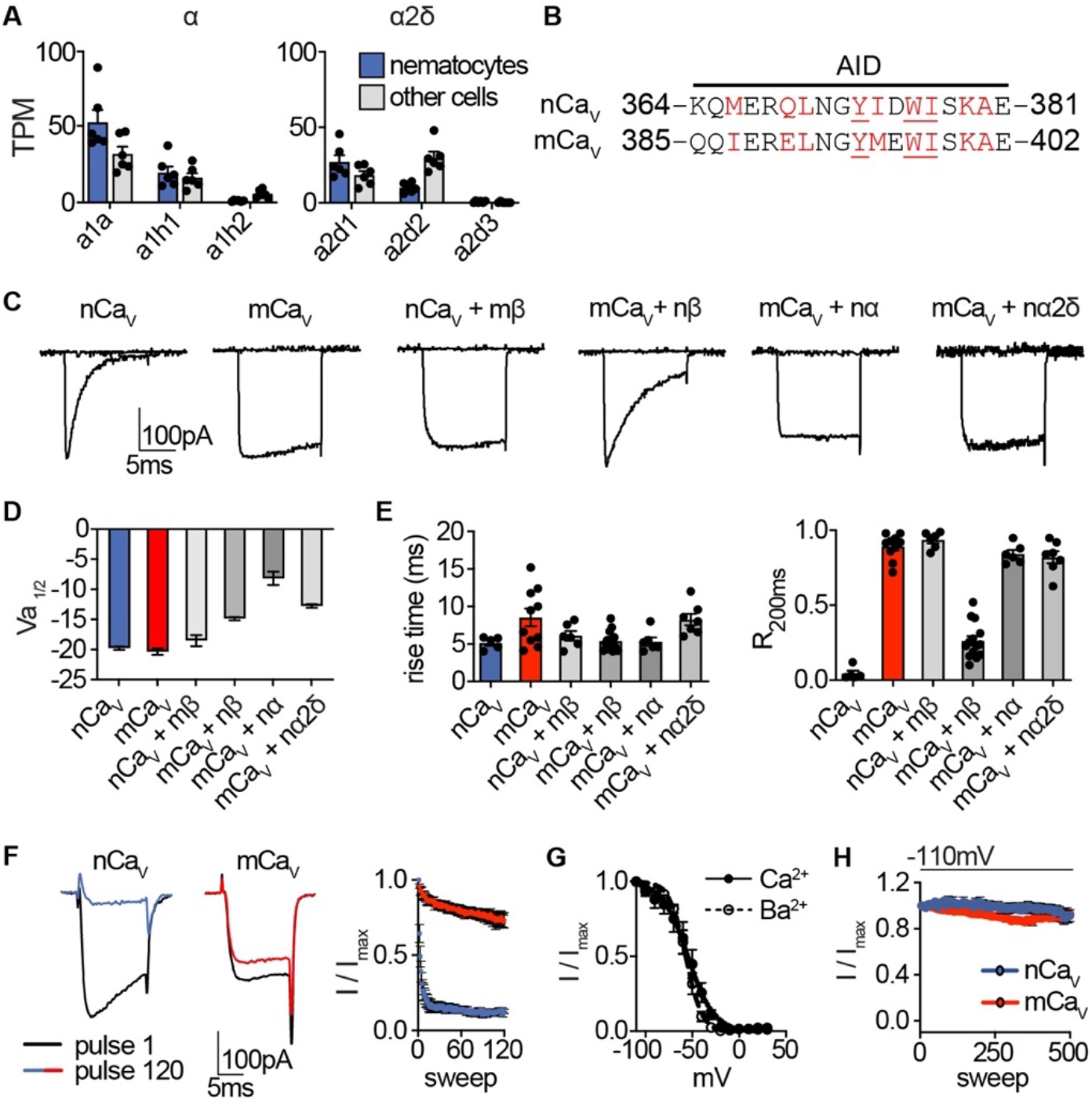
*Nematostella* Ca_v_ properties. **(A)** mRNA expression (transcripts per million, TPM) of voltage-gated calcium (Ca_v_) channel a and a2δ subunits in nematocyte-enriched cells (blue) versus non-enriched cells (grey). n = 6, p < 0.01 for cacna1a in nematocytes versus other cells, two-way ANOVA with post-hoc Bonferroni test. Expression of other subunits was not significantly different. **(B)** Sequence alignment of *Nematostella* and *Mus musculus a* subunits at the a interaction domain (AID). Red letters indicate important regions for beta subunit interaction, underlined letters are critical for interaction. **(C)** Representative voltage-activated currents from HEK-293 cells expressing wild-type or chimeric Ca_v_ channels with specific combinations of mammalian and *Nematostella* subunits. Currents were elicited by 0mV pulses from −110mV. **(D)** Half maximal activation voltage (V_a1/2_) was similar for Ca_v_ chimeras. n = 6 nCa_v_, 6 nCa_v_ + mβ, 10 mCa_v_, 10 mCa_v_ + nβ, 6 mCa_v_ + na, 6 mCa_v_ + nα2δ. **(E)** *Left:* Rise time of voltage-activated inward current. *Right:* Relative fraction of inward current remaining after a 200ms step to 0mV (R200). n = 6 nCa_v_, 6 nCa_v_ + mβ, 10 mCa_v_, 14 mCa_v_ + nβ, 6 mCa_v_ + na, 7 mCa_v_ + na2δ. **(F)** nCa_v_ inactivated more quickly than mCa_v_. Protocol: 3.33 Hz stimulation with 20ms pulses to −10mV from −90mV. n = 5 nCa_v_, 6 mCa_v_, multiple row two-tailed student’s t-test with significance of p < 0.05 by sweep 2 and p < 0.0001 by 120. **(G)** Inactivation curves of mCa_v_ + nβ were similar when external Ca^2+^ (V_i1/2_ = −54.42 ± 1.81mV, n = 10) was replaced with Ba^2+^ (V_i1/2_ = −56.95 ± 0.85mV, n = 6). **(H)** nCa_v_ exhibited relatively little slow inactivation from −110mV compared with −70mV holding potential (Fig. 2, 0.2 Hz stimulation). Data represented as mean ± sem.

**Figure 3 - figure supplement 1:**
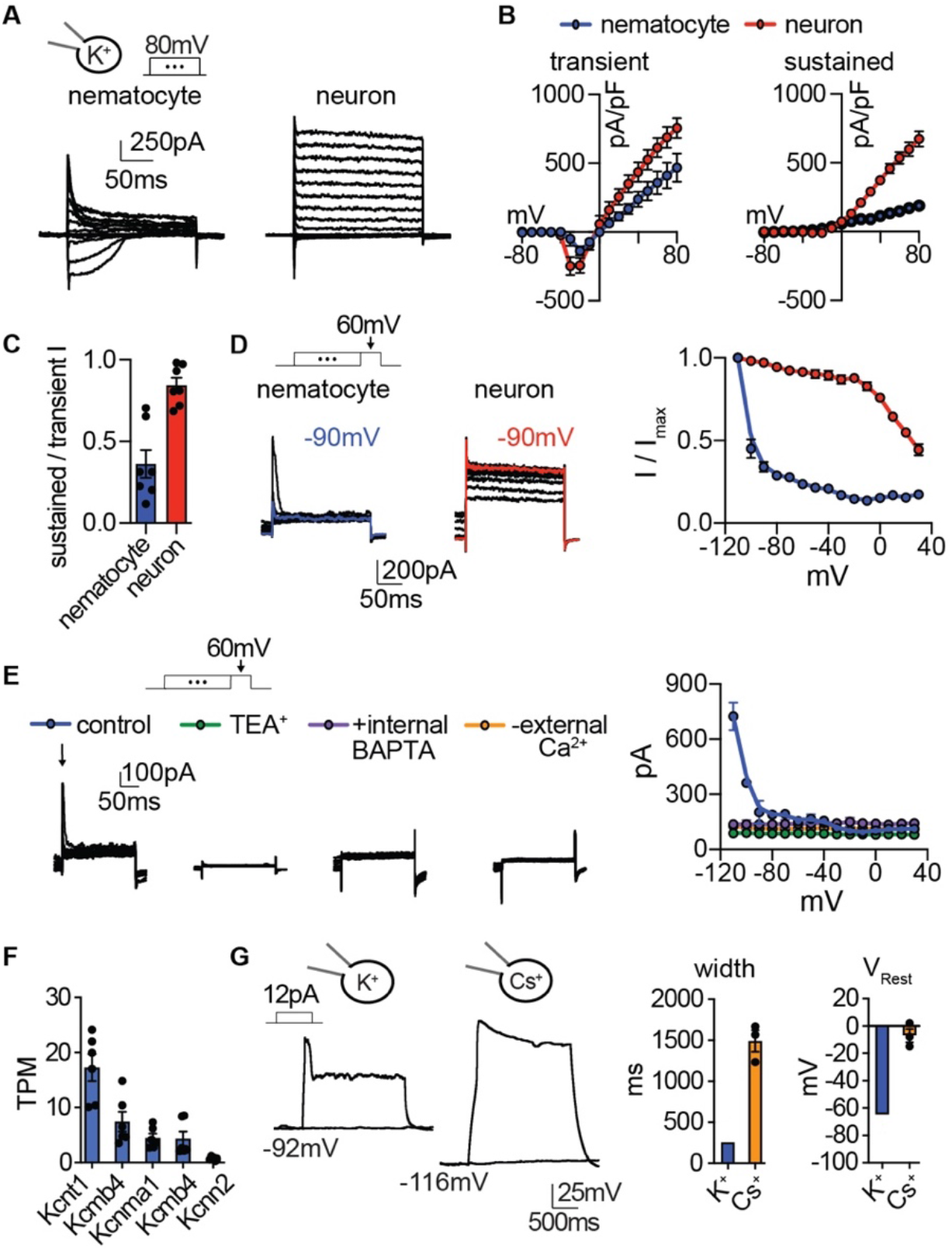
Nematocyte K^+^ current properties. **(A)** In the presence of intracellular K^+^, voltage-activated outward currents were elicited from nematocytes and tentacle neurons. **(B)** Current-voltage relationships of peak transient and sustained (end of 200ms voltage pulse) currents in nematocytes and neurons. n = 7. **(C)** Nematocytes exhibited a lower ratio of sustained / transient outward current, indicating faster inactivation of outward current. n = 7, p < 0.01 two-tailed student’s t-test. **(D)** Nematocyte transient outward currents exhibited strong voltage-dependent inactivation compared with weak voltage-dependent inactivation of outward currents in neurons. n = 5, p < 0.0001 for voltages at −100mV or above, two-way ANOVA with post-hoc Bonferroni test. **(E)** Transient K^+^ currents in nematocytes were assessed using voltage protocols to enhance the outward current sensitive to voltage-dependent inactivation. These currents were abolished by the K^+^ channel blocker TEA^+^ (n = 4), the intracellular Ca^2+^ chelator BAPTA (n = 4), or in the absence of external Ca^2+^ (replaced with NMDG^+^ + EGTA, n = 3). p < 0.0001 for currents measured following pre-pulses to −110 or −100mV, two-way ANOVA with post-hoc Bonferroni test. Smaller sustained outward currents measured following more positive pre-pulses were significantly affected by TEA^+^ but not other treatments. **(F)** mRNA expression (transcripts per million, TPM) of Ca^2+^-activated K^+^ channels in nematocyte- enriched cells. n=6. **(G)** Nematocyte spike width and resting membrane potential were affected by the presence of intracellular Cs^+^. Cells were first hyperpolarized to elicit spikes with subsequent current injection. Spike width: n = 8 K^+^ (from Fig. 3) and 3 Cs^+^, p < 0.05 two-tailed student’s t-test. Resting membrane voltage: n = 12 K^+^ (from Fig. 3) and 3 Cs^+^, p < 0.01 two-tailed student’s t-test. Data represented as mean ± sem.

**Figure 4 - figure supplement 1:**
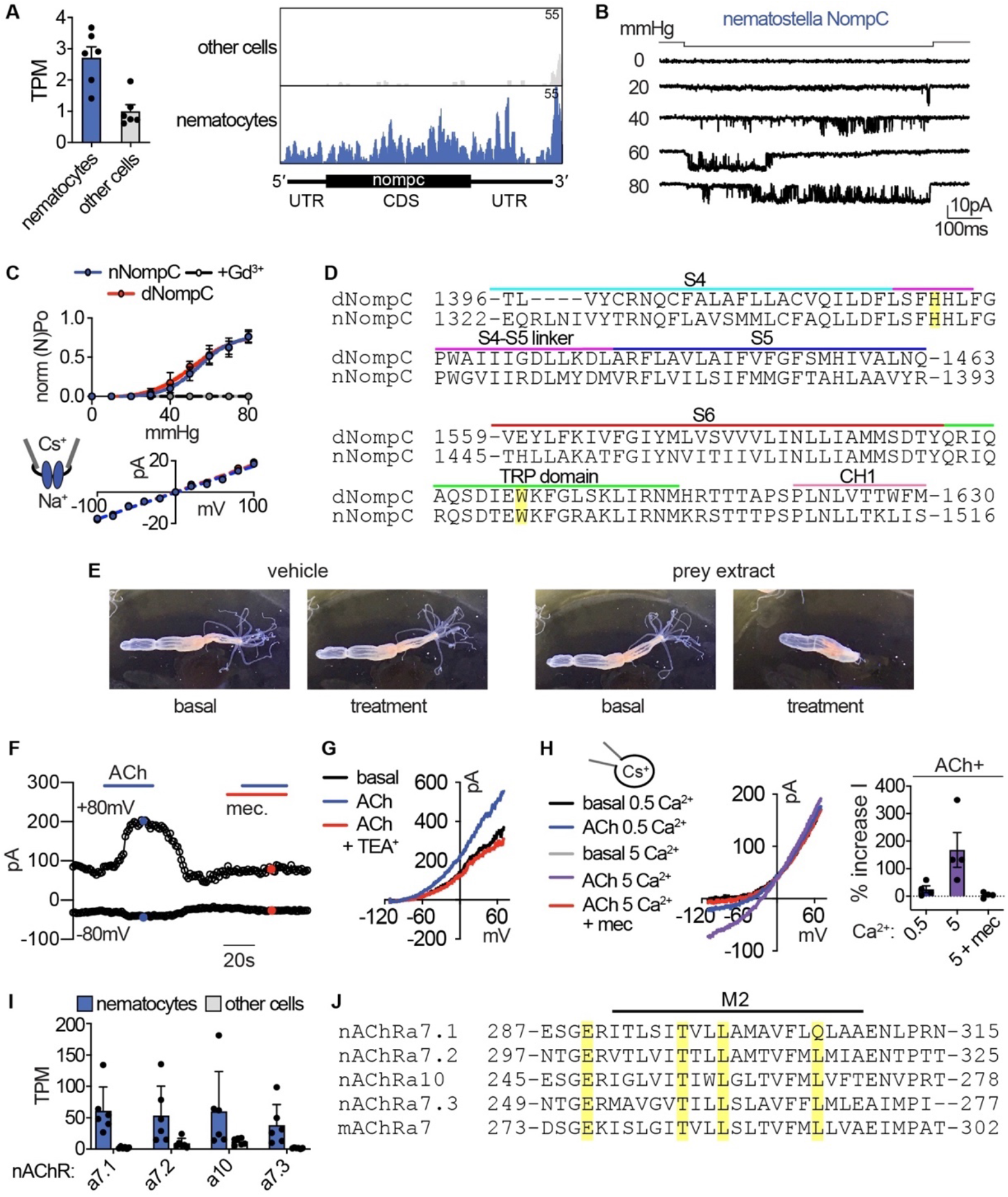
Sensory transduction properties. **(A)** *Left:* mRNA expression (transcripts per million, TPM) of NompC (no mechanoreceptor potential) ion channels in nematocyte-enriched cells (blue) vs. nonenriched cells (grey). n = 6, p < 0.05, two-tailed student’s t-test. *Right:* Numerous reads mapped across the entire NompC sequence. The transcript architecture is shown below the mapping. CDS, coding sequence; UTR, untranslated regions. The maximum read counts was set as the same value between samples. **(B)** A representative single *Nematostella* NompC (nNompC) channel expressed in an excised membrane patch was sensitive to increasing pressure applied via the patch pipette. Vm = −80mV. **(C)** nNompC and *Drosophila* NompC (dNompC) exhibited similar pressure-open probability (P_O_) relationships and slope conductance. nNompC was blocked by Gd^3+^. 95% confidence interval for pressure required to induce half maximal P_O_: nNompC = 51.85 – 59.44 mmHg, dNompC = 47.44 – 59.73 mmHg. n = 5. Slope conductance: nNompC = 168.3 ± 4 pS, dNompC = 174.8 ± 4 pS. n = 4. **(D)** Alignment of *Drosophila* (dNompC) and *Nematostella* (nNompC) protein sequences revealed high conservation with overall sequence identity of 44.3% (62.7% similarity). Yellow indicates residues important for mechanosensitivity. **(E)** Application of filtered prey extract, but not vehicle, elicited a feeding response (contraction of tentacles) in a representative *Nematostella.* **(F)** Representative nematocyte response to acetylcholine (ACh) shows an outwardly- rectifying current, which was blocked by mecamylamine (Mec). K^+^ was the major intracellular cation. **(G)** Representative current-voltage relationship shows that the ACh-evoked current was inhibited by the K^+^ channel blocker TEA^+^. **(H)** In the presence of intracellular Cs^+^ to block K^+^ currents, ACh mediated a mecamylamine-sensitive, inward current that was enhanced by extracellular Ca^2+^. **(I)** mRNA expression (transcripts per million, TPM) of nicotinic acetylcholine-like receptors (nAChRs) in nematocyte-enriched cells (blue) vs. non-enriched cells (grey). n = 6, p < 0.05 for nACHRa7.1, two-way ANOVA with post-hoc Bonferroni test. **(J)** Alignment of *Nematostella* nAChRs and mouse nAChRa7 protein sequences revealed conserved residues that facilitate Ca^2+^ permeability. Data represented as mean ± sem.

**Figure 5 - figure supplement 1:**
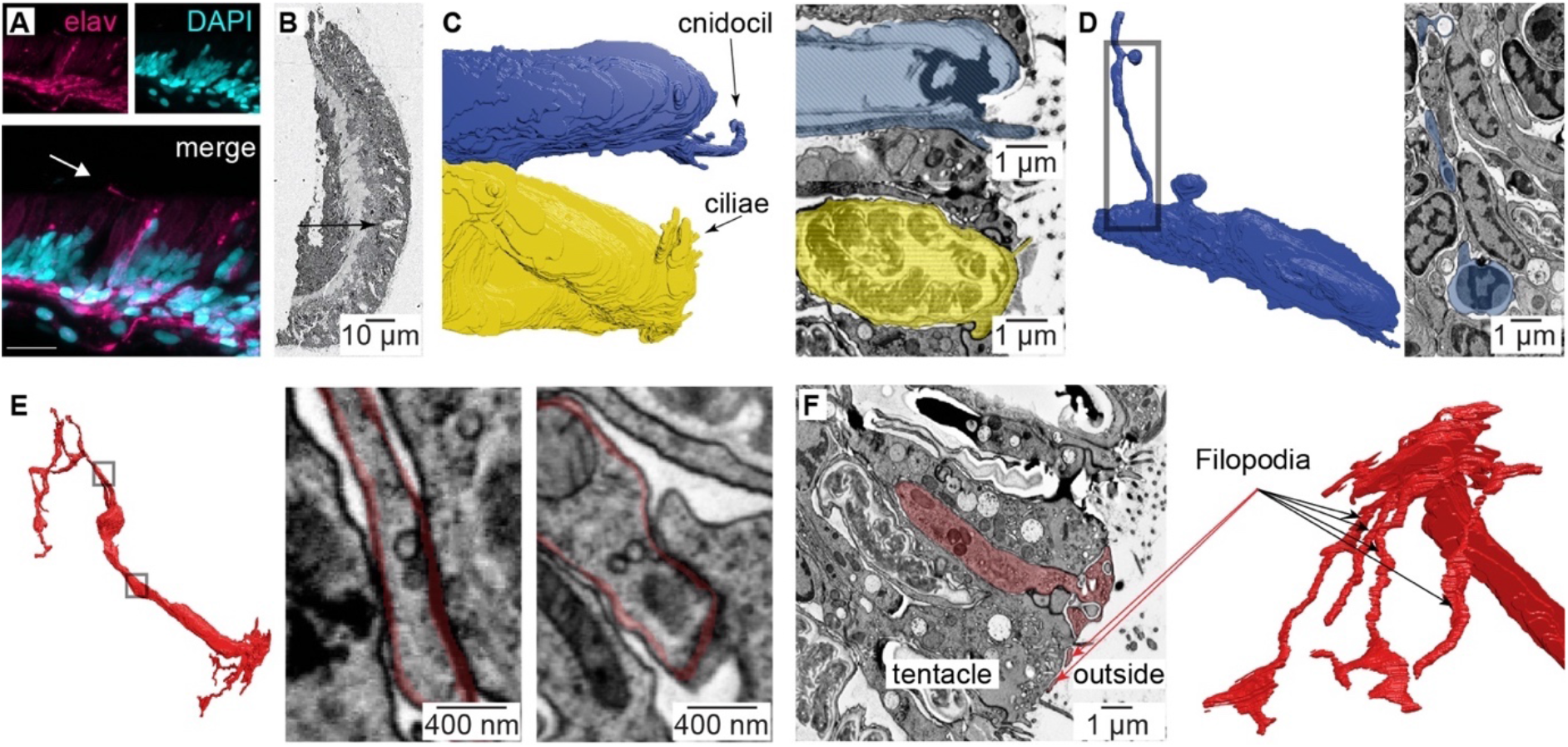
Nematocyte morphology. **(A)** Tentacle from elav::mOrange *Nematostella* stained with dsRed (red) indicated neural processes beneath ectodermal nematocytes and putative sensory neurons located within the ectoderm. DAPI (blue) stains nuclei and nematocysts. White arrow indicates putative sensory projection. Scale bar = 10μm. **(B)** Electron micrograph of *Nematostella* tentacle section (50nm thickness). Arrow indicates regions containing nematocytes, spirocytes and sensory neurons which were imaged at higher magnification as illustrated in Fig. 5 and Fig. 5 - supplement 2. **(C)** Cnidocil of a nematocyte (blue) and ciliae of a spirocyte (orange). *Left:* 3D reconstruction. *Right:* Electron micrograph of one section of the nematocyte (*top*) and one section of the spirocyte (*bottom*). **(D)** Nematocyte process. *Left:* 3D reconstruction of the nematocyte. *Right:* Electron micrograph of one section across the nematocyte process, indicated by the region boxed on the left panel. **(E)** Synapses made between the neuron and other cells than a nematocyte. Two examples are shown (middle and right panels). **(F)** Sensory terminal of the neuron shown in Fig. 5A with multiple filopodia extending in the outside environment. *Left:* Electron micrograph of one section of the sensory neuron. *Right:* 3D reconstruction of the sensory neuron.

**Figure 1 - figure supplement 2:**
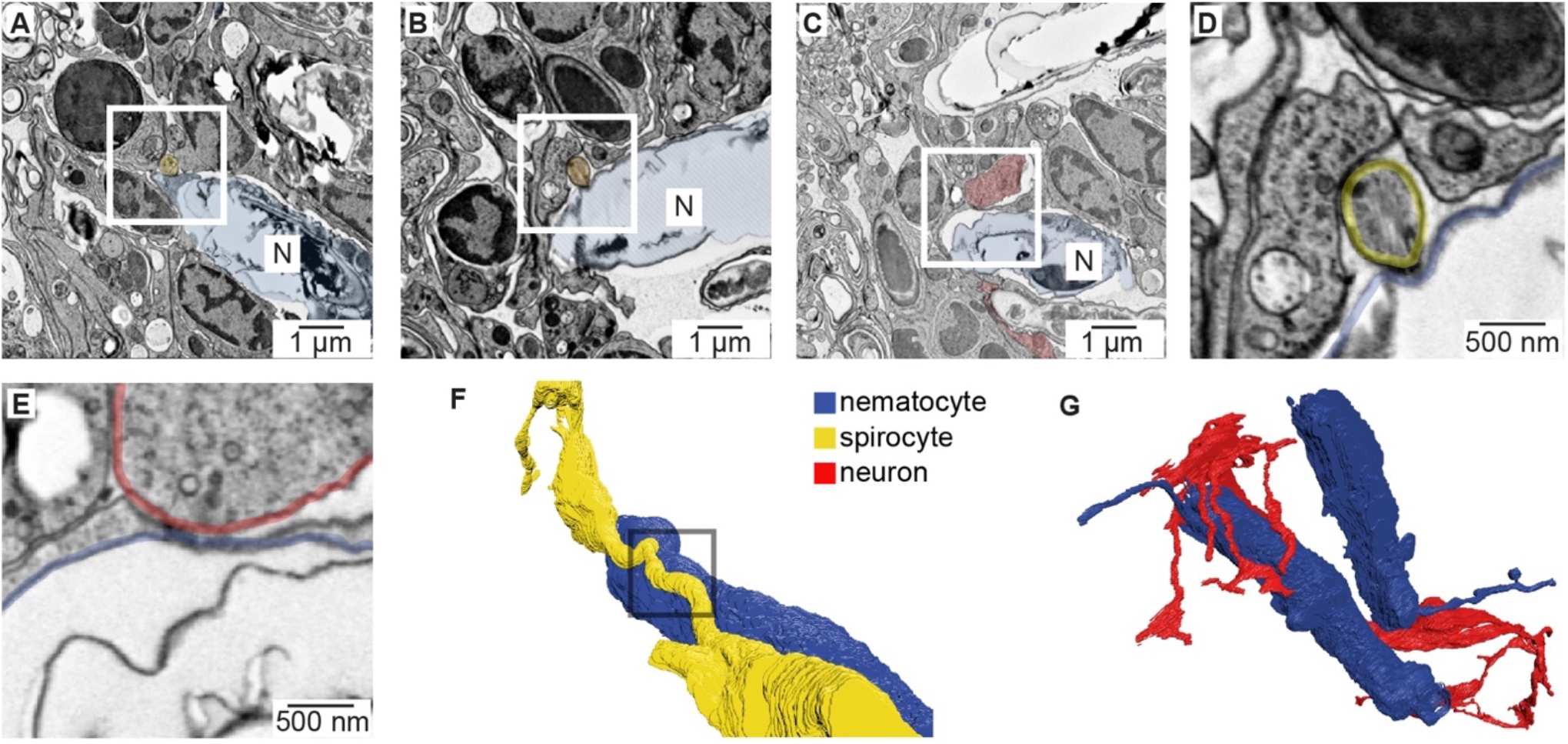
Nematocyte synaptic connections. **(A)** Putative synapse between a nematocyte (N, blue) and a spirocyte (yellow). Higher magnification is shown in Fig. 5A. **(B)** Putative synapse between a nematocyte (N, blue) and a spirocyte (orange). **(C)** Putative synapse between a nematocyte (N, blue) and a sensory neuron (red). **(D)** Higher magnification of the regions boxed in B. **(E)** Higher magnification of the regions boxed in C. **(F)** Reconstruction of the synapse between a nematocyte (N, blue) and a spirocyte (S, yellow). Box indicates site of synaptic contact shown in panel A and Fig. 5A. **(G)** 3D reconstruction of the sensory neuron and the two nematocytes shown in Fig. 5A (blue).

**Figure 5 - figure supplement 3:**
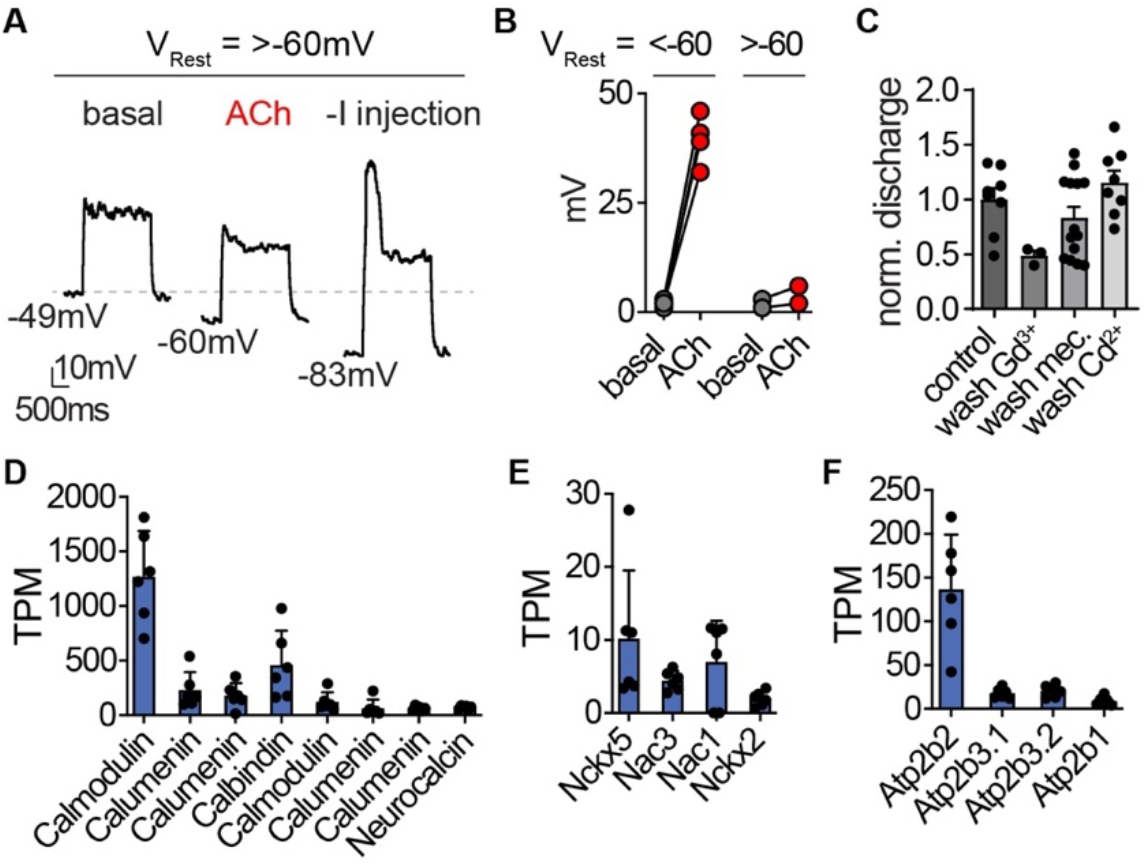
Nematocyte signaling pathways. **(A)** For nematocytes with a resting membrane voltage (V_Rest_) more positive than −60mV, ACh was insufficient to hyperpolarize cells enough to mediate a spike following depolarizing current injection. Representative of n = 2. **(B)** For nematocytes with V_Rest_ more negative than - 60mV, all ACh-hyperpolarized cells produced voltage spikes following depolarizing current injection. n = 4, p < 0.001 two-tailed student’s t-test. **(C)** Controls (related to Figs. 1G and 5D) for nematocyte discharge. Control = 8, Gd^3+^ wash = 7, Mec wash = 14, Cd^2+^ wash = 8. **(D-F).** mRNA expression (transcripts per million, TPM) of calcium-binding proteins (**D**), calcium exchangers (**E**), and calcium ATPases (**F**) in nematocyte-enriched cells.

